# Neratinib, a clinical drug against breast cancer, protects against atherosclerosis via ASK1 inhibition

**DOI:** 10.1101/2024.11.05.622073

**Authors:** Fanshun Zhang, Yanjun Yin, Zhihua Wang, Xiumei Wu, Danielle Kamato, Jianping Weng, Suowen Xu

## Abstract

Atherosclerosis commences with endothelial dysfunction and the retention of cholesterol within the vessel wall, followed by a chronic inflammatory response. Cholesterol-lowering strategies (such as statins and PCSK9 inhibitors) are primarily used for treating patients with atherosclerotic cardiovascular diseases, but leaving the therapeutic dilemma of residual inflammatory risk. To address this challenge, we employed Connectivity Map (CMap) screening for inflammation mechanism-based anti-atherosclerotic compounds using perturbational datasets obtained from TNFα and IL-1β-stimulated human endothelial cells. This screening process allow us to identify Neratinib, a clinical drug against breast cancer, as the hit compound with potential anti-inflammatory actions in endothelial cells. Further studies reveal that Neratinib inhibited endothelial cell inflammation elicited by three different pro-inflammatory stimuli (TNFα, IL-1β and LPS). Intriguingly, the anti-inflammatory effect of Neratinib was independent of its classical target HER2/ERBB2 inhibition. Mechanistically, Neratinib directly binds ASK1 and suppresses ASK1 activation. In both male and female *Ldlr*^−/−^ mice, treatment with Neratinib decreased the plaque area, reduced the necrotic core size and mitigated macrophage infiltration to stabilize plaques. Lastly, we observed that Neratinib, in conjunction with the use of Rosuvastatin (a standard lipid-lowering drug), led to a reduction in serum lipids, and produced synergistic anti-atherosclerotic effects. Olink proteomics study suggested that combination treatment alleviated inflammation-related cytokines/chemokines in the serum from *Ldlr*^−/−^ mice. Taken together, these findings support the concept that Neratinib could be tested for its potential as a “repurposed” drug for vascular inflammation and atherosclerosis, thereby streamlining efforts to translate preclinical discoveries to clinical testing in humans.

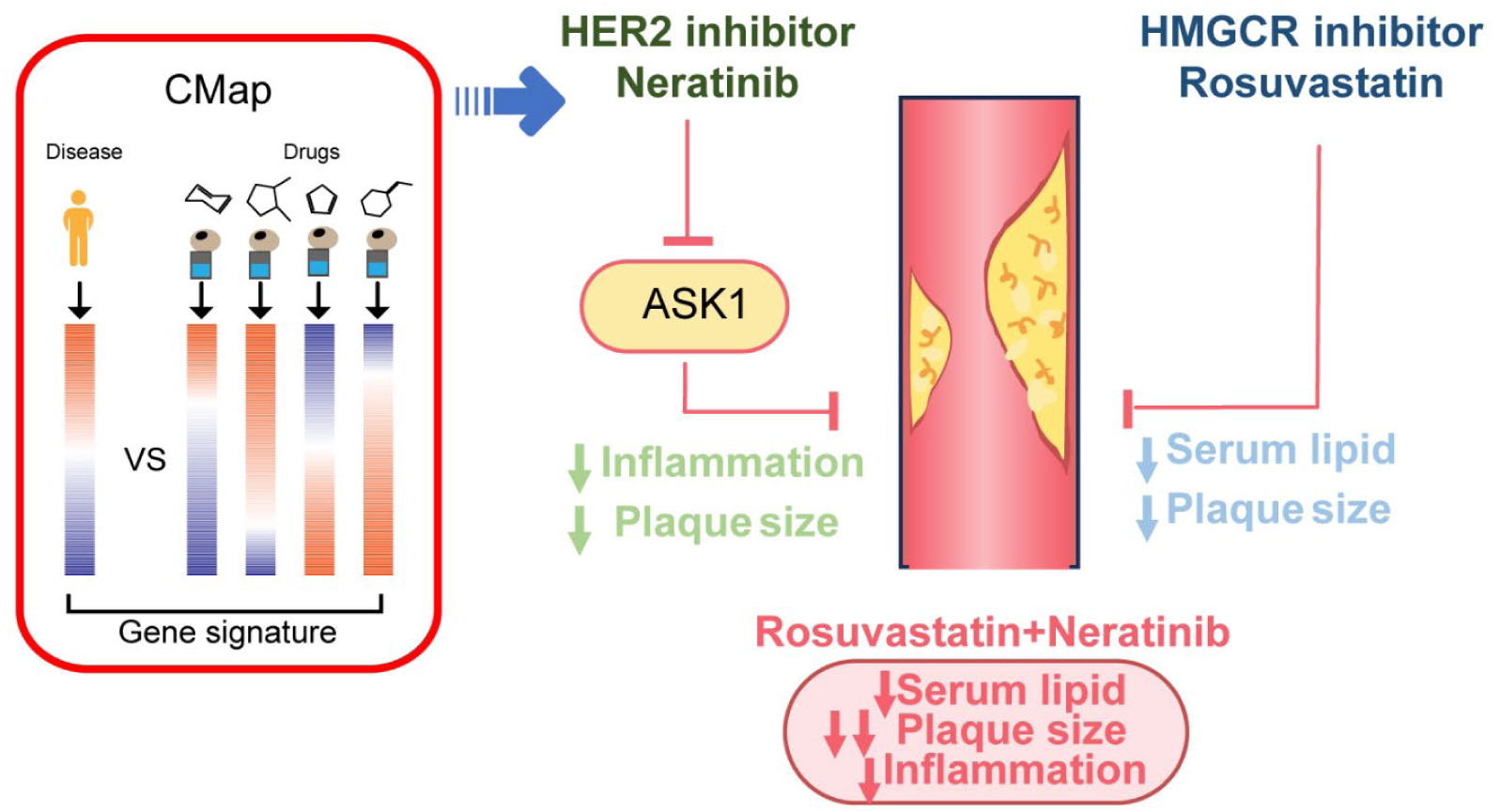

**One Sentence Summary:** Neratinib protects against atherosclerosis by reducing endothelial inflammation via ASK1 inhibition

## INTRODUCTION

Atherosclerotic cardiovascular disease (ASCVD) remains the leading cause of morbidity and mortality globally. ASCVD is a chronic inflammatory disease triggered by the accumulation of cholesterol-rich lipoproteins in the vessel wall. ASCVD commences with the retention of low-density lipoproteins (LDL) within the vessel wall, while retained LDL are susceptible to oxidation, thus forming the notorious oxidized LDL (oxLDL). OxLDL are engulfed by macrophages and leads to the formation of foam cells, the hallmark of atherosclerosis. Atherosclerotic plaques can become “unstable” as a result of unresolved inflammation, erosion and rupture leading to myocardial infarction and stroke^1, 2^. The current gold standard treatment for ASCVD is to reduce LDL levels with HMG-CoA reductase inhibitors (e.g., Statins), PCSK9 inhibitors and many others.

Beyond a lipid-driven disease, ASCVD is also recognized as a chronic inflammatory disease^2^. In the development of atherosclerosis, systemic risk factors (such as oxLDL, hyperglycemia etc) contributes to the increased secretion of inflammatory factors (TNF-α, IL-1β, and IL-6 etc) from vascular cells and creates a local inflammatory microenvironment which propagate the pro-atherogenic cascade. Emerging evidence from preclinical and clinical studies have consolidated that anti-inflammatory therapy is effective in reducing major adverse cardiovascular events. At present, the cardiovascular protective actions of anti-inflammatory therapies, such as Colchicine^3, 4^, Canakinumab (an IL-1β monoclonal antibody)^3, 5^, Ziltivekimab (an IL-6 monoclonal antibody)^6^ have been well documented. However, the number of drugs with anti-inflammatory effects for the treatment of atherosclerosis is still very limited (only colchicine is approved by FDA). Also, some anti-inflammatory therapies impair host’s immune response. For example, the use of IL-1β targeted therapy Canakinumab was associated with a higher incidence of fatal infection^3^. Last but not least, high attrition rates, substantial costs and slow pace of traditional drug discovery have hampered new drug discovery and development efforts^7^, Therefore, there is unmet medical need to identify inflammation mechanism-based drugs which can attenuate atherosclerosis development effectively and safely. To this end, repurposing of FDA-approved drugs to treat atherosclerosis is increasingly becoming an attractive proposition due to potentially lower-risk profile, lower costs and shorter development timelines ^7^. Considering the dual acting mechanisms of atherosclerosis (dyslipidemia and inflammation), anti-inflammatory drugs can be posited as important complementary therapies for the treatment of ASCVD^8^.

Connectivity map (CMap) is a dataset of genome-wide expression profile from cells that have been exposed to pharmacological or genetic perturbagens^9^ thereby connecting the bridge between drugs, genes and disease^10^. Based on the principle that drugs with similar genome-wide expression signatures most possibly share same direction of effect for diseases, CMap is thus a systematic approach for drug repurposing. Here, we employed in *silico* CMap exploration and cell experiments to identify hit compounds that target endothelial inflammation. Our screening process identified Neratinib, an orally bioavailable small-molecule tyrosine kinase inhibitor for treating patients with early-stage HER2^+^ breast cancer^11^, as a novel anti-inflammatory drug. Neratinib significantly reduced endothelial inflammatory response via ASK1 inhibition and protects against atherosclerosis through alleviating systemic inflammation. Further, administrating Neratinib concurrently with Rosuvastatin achieved synergistic effects in attenuating atherosclerosis development. Taken together, this study offered a potentially new therapeutic drug to reduce residual cardiovascular risk for patients with atherosclerosis by drug repurposing.

## RESULTS

### Discovery of Neratinib as a novel anti-inflammatory drug based on CMap screening

To identify the shared inflammatory gene signature involved in endothelial cell inflammation, human umbilical vein endothelial cells (HUVECs) were stimulated with inflammatory stimuli TNF-α or IL-1β, two prototypic stimuli involved in triggering inflammatory responses in endothelial cells. RNA-sequencing transcriptomic data were analyzed to select top 150 upregulated and downregulated genes with the criteria of *q* value < 0.05 and log_2_(FC) > 1 or log_2_(FC) < −1 (Supplement Table 1 and Supplement Table 2). The top 150 genes with significant upregulation or downregulation were constructed as the inflammation dataset (Figure. 1A), which were uploaded to the CMap database, matched and scored to obtain drug perturbed signature in inverse relation to that of TNF-α and IL-1β (Supplement Table 3). We defined the parameter “S” to represent the score of the compound in lowering TNF-α and IL-1β induced gene signature (Figure. 1B). According to the “S” ranking value (high, medium and low), three groups of drugs, with a total of 24 compounds, were used for verification of anti-inflammatory effects in cultured endothelial cells (Supplement Table 3). The expression of ICAM1 and VCAM1 in the presence of TNF-α or IL-1β was selected as the functional readout. The anti-inflammatory effects of the compounds were evaluated by the average ranks (*R̅*) of all the evaluated items. A smaller average rank reflected a stronger anti-inflammatory effect. The results of *R̅* yield Neratinib as a leading anti-inflammatory agent (drug No. 12), followed by Copanlisib (drug No. 3), Ponatinib (drug No. 4) and Cilengitide (drug No. 9) (Figure. 1C and 1D). The potential anti-atherosclerotic and anti-inflammatory effects of Copanlisib^12^, Ponatinib^13, 14^ and Cilengitide ^15^ have been reported, which testified the efficacy of our screening model. Considering that side effects also were observed in the literature, the three drugs were not selected for further analysis ^16–18^. Since there are no reports regarding the actions of Neratinib in endothelial cells and atherosclerosis, Neratinib was selected for further evaluations.

**Figure 1.**
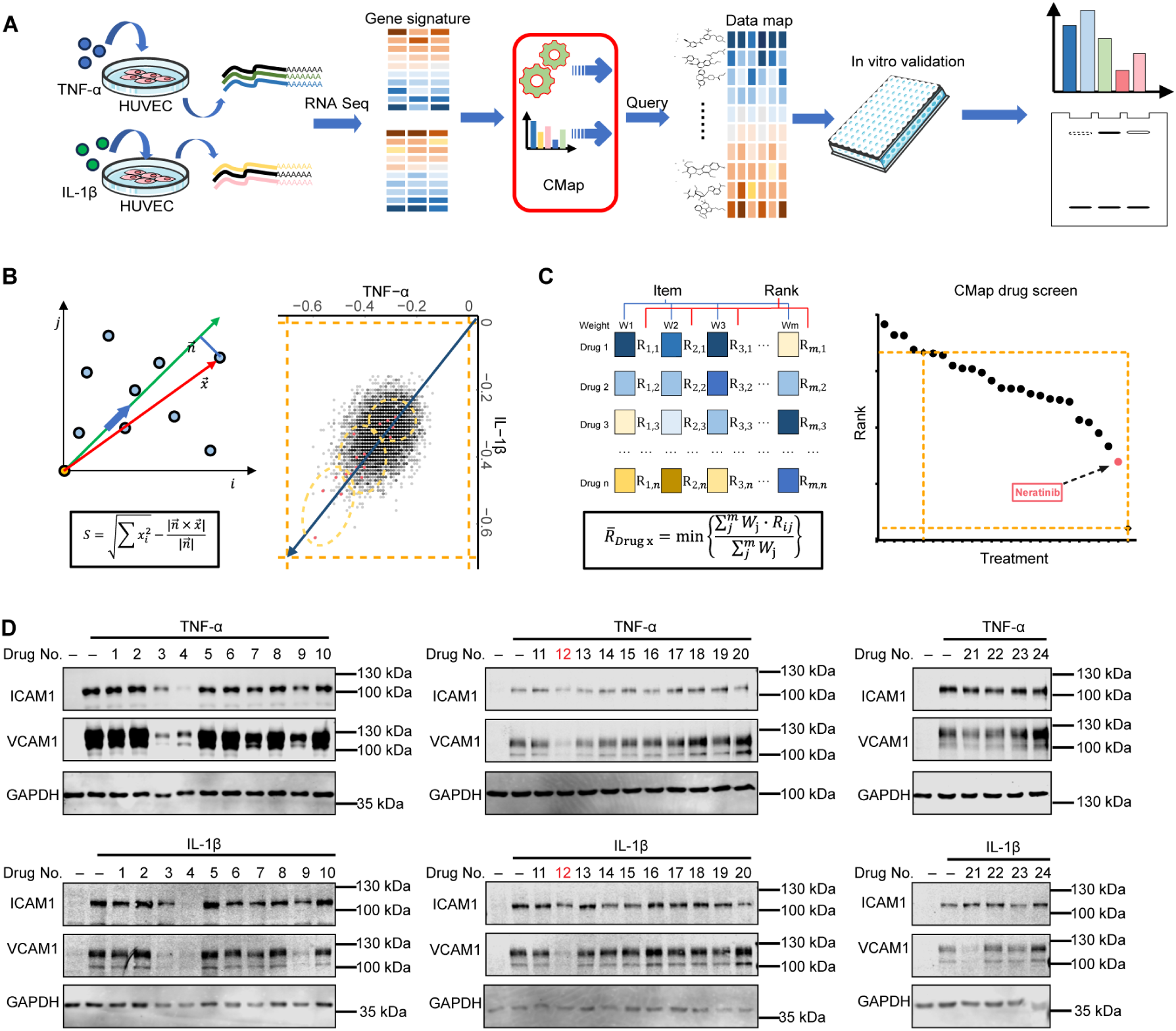
Discovery of Neratinib as a novel anti-inflammatory drug based on CMap screening. (A) Scheme of CMap-screening. (B) Algorism for the CMap-based screening. (C) Experimental verification. At the cellular level, ICAM1 and VCAM1 expression in endothelial cells stimulated with TNF-α and IL-1β were selected as screening readout. The anti-inflammatory effects of the compounds were evaluated by the average ranks (*R̅*) of all the evaluated items. A smaller average rank reflects a stronger anti-inflammatory effect. (D) Western blot verification of identified drugs which inhibit endothelial inflammation. HUVECs are treated with indicated drugs at 5 μM for 24 hours and then stimulated with 10 ng/ml TNF-α or IL-1 β for 6 hours; n=3 biologically independent repeats for each drug. Red, compound 12 (Neratinib).

We next investigated the anti-inflammatory effects of Neratinib in HUVEC. Endothelial cells responded to the stimulation of TNF-α, characterized by the upregulation of inflammatory genes (*IL-6*, *CCL2*, *ICAM1*, *VCAM1*, and *SELE*). This inflammatory phenotype was suppressed by treatment with Neratinib (Figure. 2A). A similar response was observed when endothelial cells were treated with Neratinib in the presence of inflammatory stimuli IL-1β or LPS (Figure. 2B and 2C), suggesting the universality of Neratinib-mediated anti-inflammatory effects. Neratinib reduced TNF-α or IL-1β-induced elevated protein levels of ICAM1 and VCAM1 in a concentration-dependent manner as well as THP-1 adhesion to inflamed human endothelial cells (Figure. 2D-2F). The anti-inflammatory effects of Neratinib were also recapitulated in human aortic endothelial cells (HAEC) (Figure. 2G and 2H). These results demonstrated the anti-inflammatory actions of Neratinib in human endothelial cells.

**Figure 2.**
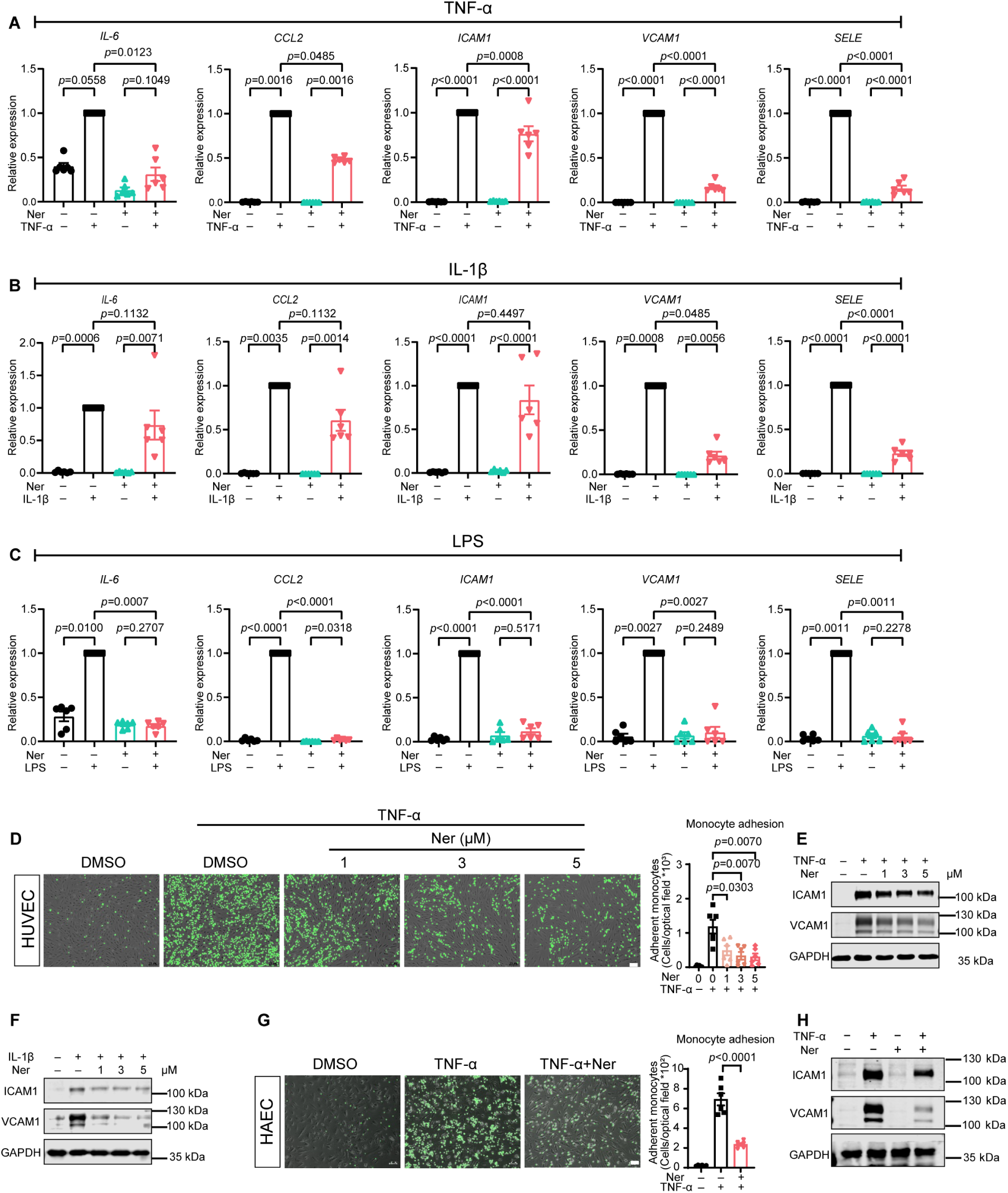
Neratinib alleviates endothelial cell inflammation. (A-C) The expression of inflammatory genes in HUVECs exposed to various pro-inflammatory treatments. HUVECs were pretreated with 5 μM Neratinib before stimulation with 10 ng/ml TNF-α, 10 ng/ml IL-1β or 1 μg/ml LPS; n=6 biological independent repeat for each group. (D) THP1 monocyte adhesion assay (in HUVECs); n=6 biological independent repeats for each group. Scale bar: 20 μm. (E and F) Western blot results of HUVECs stimulated with 10 ng/ml TNF-α or 10 ng/ml IL-1β to show the protein levels of ICAM1 and VCAM1; n=6 biological independent repeats for E and n=3 for F. (G) THP1 monocyte adhesion assay (in HAECs); n=6 biological independent repeats for each group. Scale bar: 20 μm. (H) The protein levels of ICAM1 and VCAM1 in HAECs treated with 5 μM Neratinib for 30 minutes and stimulated with 10 ng/ml TNF-α; n=7 biological independent repeats for each group. All data are presented as the means ± SEMs. Data of *IL-6* and *CCL2* in A, *IL-6*, *CCL2* and *VCAM1* in B and *IL-6*, *VCAM1* and *SELE* in C are analyzed using Kruskal-Wallis test. Data of *ICAM1*, *VCAM1* and *SELE* in A, *ICAM1* and *SELE* in B and *CCL2* and *ICAM1* in C are analyzed using one-way ANOVA test. Data of D are analysed using Kruskal-Wallis test. Data of G are analyzed using Brown-Forsythe and Welch ANOVA test.

### Neratinib protects against atherosclerosis in *Ldlr*^−/−^ mice

Based on the anti-inflammatory effects of Neratinib, we reasoned whether Neratinib can prevent atherosclerosis development in mice. We first explored the safe doses of Neratinib (Supplement Figure. 1A). Normal male C57BL/6J mice were administered with 20 mg/kg/day and 60 mg/kg/day Neratinib via oral gavage and we found that Neratinib does not affect body weight, organ index, serum biochemistry parameters or tissue damage except that high-dose Neratinib increased serum urea levels (Supplement Figure. 1B and 1G). Therefore, we selected the dose of 20 mg/kg/day for our pharmacological intervention study in *Ldlr*^−/−^ mice fed a high-cholesterol diet for 14 weeks (Figure. 3A). We observed no difference in body weight, serum lipid levels, serum markers associated with tissue injury or fasting blood glucose levels between Neratinib- and vehicle-treated mice (Supplement Figure 2). To assess the effect of Neratinib on atherosclerosis progression and plaque composition, we analyzed plaque area in mouse aorta and aortic sinus, necrotic core size, collagen content, and CD68- and α-SMA-positive areas. In Neratinib-treated group, fewer plaques, as shown by Oil Red O staining, are observed in the *en face* aorta and aortic sinus in both male and female mice (Figure. 3B and 3C, Supplement Figure. 3A and 3B). HE staining revealed that the size of necrotic core within the aortic sinus in the Neratinib treatment group was significantly smaller than that in the control group (Figure. 3D and Supplement Figure. 3C). Neratinib did not affect the collagen content in the plaque from either male or female *Ldlr*^−/−^ mice (Figure. 3E and Supplement Figure. 3D). We also examined macrophage infiltration and smooth muscle cells content via immunofluorescence staining. The results revealed a significant decrease in macrophage infiltration in male mice and an increase in the content of smooth muscle overlying the plaques after neratinib treatment (Figure. 3F and Supplement Figure. 3E). These data indicated that the administration of Neratinib significantly protected against atherosclerosis.

**Figure 3.**
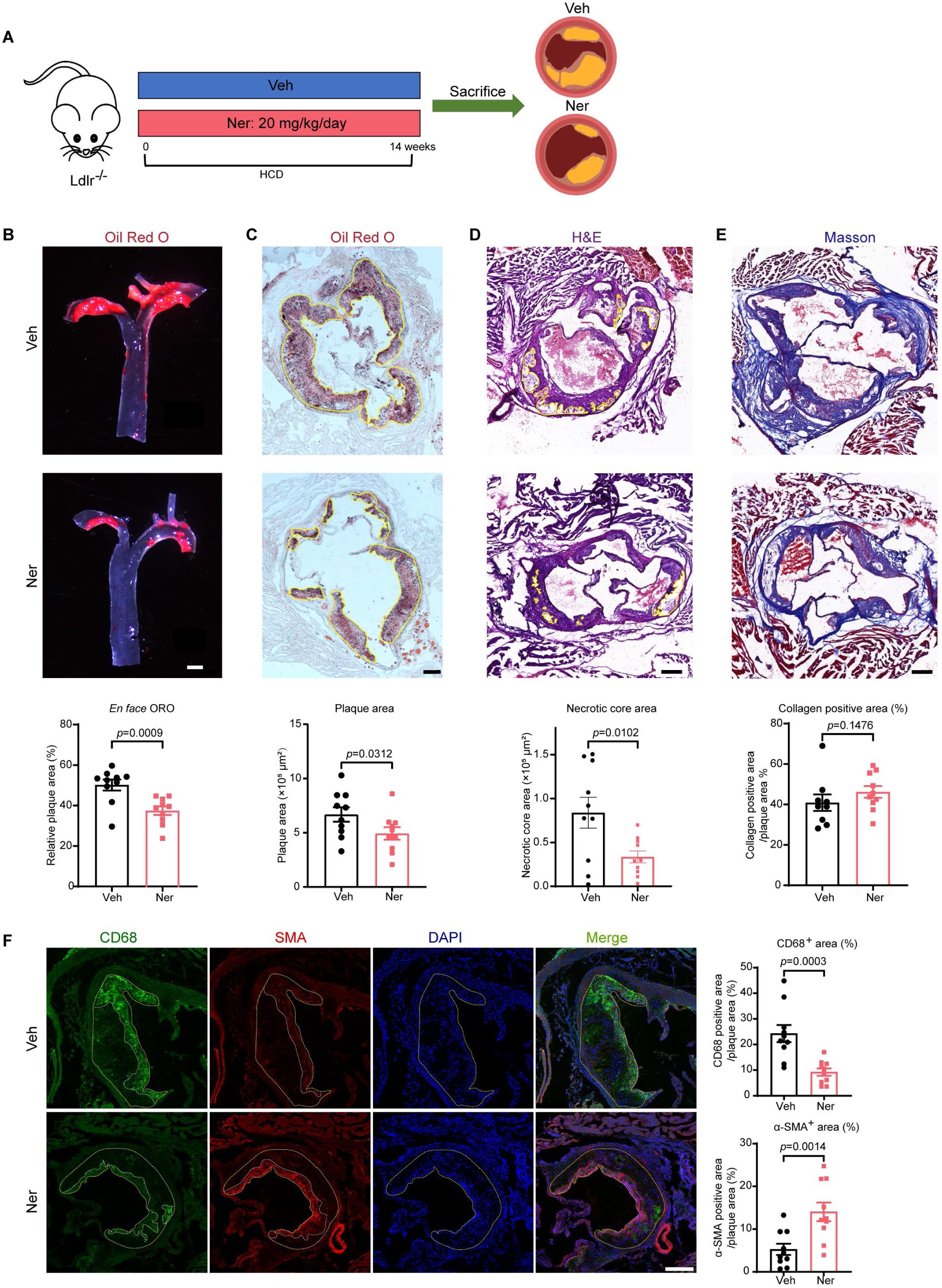
Neratinib protected against atherosclerosis in *Ldlr*^−/−^ mice. (A) Flow chart of the animal experiment. *Ldlr*^−/−^ mice were fed a high-cholesterol diet (HCD) for 14 weeks and were concurrently administered with 20 mg/kg Neratinib by oral gavage. (B) Oil Red O staining of the *en face* aorta. Scale bar: 1 mm, n=10 for each group. (C) Oil Red O staining of aortic sinus. Scale bar: 200 μm, n=10 for each group. (C and D) H&E and Masson staining of aortic sinus cryosections. Necrotic core size and collagen content in plaques was quantified. Scale bar: 200 μm, n=10 for each group in C and n=9 for vehicle group and n=10 for Ner group in D. (F) Immunofluorescence staining of CD68 and α-SMA in the aortic sinus. CD68 positive infiltrated macrophages and α-SMA positive smooth muscle cell content were quantified. Scale bar: 200 μm, n=10 for each group. All data are presented as the means ± SEMs. Data of B are analyzed using Kolmogorov-Smirnov test. Data of C, E and α-SMA^+^ area in F were analyzed using unpaired Student’s t-test. Data of D and CD68^+^ area in F were analysis using unpaired Student’s t-test with Welch’s correction.

### Neratinib attenuates endothelial cell inflammation independent of HER2/ERBB2 inhibition

Neratinib is a small-molecule tyrosine kinase inhibitor that irreversibly inhibits the activation of human epidermal growth factors (HER1/EGFR, HER2/ERBB2, and HER4/ERBB4). HER2, encoded by *ERBB2*, is the classical target of Neratinib. To profile the expression of HER family members in the vasculature, we analyzed publicly deposited single-cell RNA-seq from mouse aorta and human carotid artery. The results show the expression of all members of HER family maintained a low level of expression in both mouse and human arterial endothelium using *VWF* as the positive control for endothelial cell-enriched gene (Supplement Figure. 4A and 4B). To explore whether the expression of HER family members is altered by inflammation, we analyzed the transcriptome sequencing results of TNF-α-stimulated HUVECs. All members of the *ERBB* family remained at low levels, and their expression was not influenced by TNF-α treatment (Supplement Figure. 4C). *HER2*/*ERBB2* is a key member in the *ERBB* family, and the fpkm value of *ERBB2* is the highest among those of the *ERBB* family. We next investigated whether pharmacological or genetic inhibition of the ERBB2 signaling pathway could achieve anti-inflammatory effects. Unexpectedly, silencing of ERBB2 with siRNA did not reduce TNF-α-stimulated ICAM1 or VCAM1 expression at protein levels. Also, Neratinib exhibits comparable anti-inflammatory effects in *ERBB2*-depleted endothelial cells (Supplement Figure. 4D). In addition, we used Pyrotinib^19, 20^ and Lapatinib^21, 22^ (two widely used EGFR/HER2 dual inhibitors in clinical trials) to inhibit ERBB2 activity. Both Pyrotinib and Lapatinib significantly inhibit the phosphorylation of ERBB2 but did not reduce the expression of ICAM1 or VCAM1 (Supplement Figure. 4D and 4F). Additionally, the anti-inflammatory effect of Neratinib persists when *ERBB2* was overexpressed in HUVECs by adenovirus (Supplement Figure. 4G). The results of dual-luciferase reporter assay show ERBB2 overexpression inhibits NF-κB transcriptional activity and did not affect the inhibition effect of Neratinib (Supplement Figure. 4H). These results suggest that the anti-inflammatory effect of Neratinib is independent of ERBB2 inhibition.

**Figure 4.**
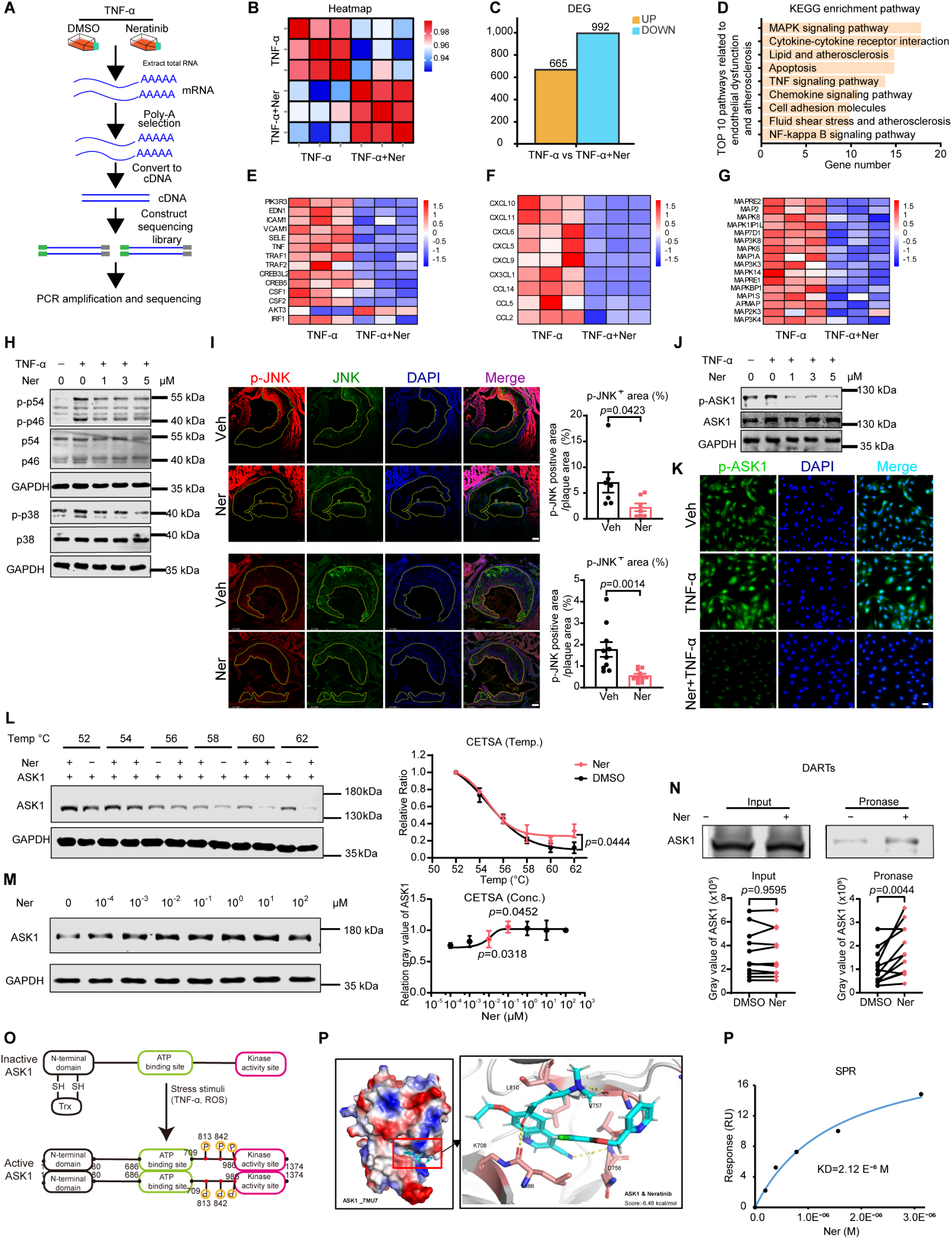
Identification of ASK1 as the molecular target of Neratinib. (A) Scheme of RNA-sequencing; n=3 biological independent samples for each group. (B) Heatmap of sequencing. (C) Differentially expressed genes (DEGs) in the TNF-α + Ner vs TNF-α groups. (D) KEGG enrichment pathway analysis of down-regulated genes related to endothelial dysfunction and atherosclerosis. MAPK pathway ranks the top. (E and F) Heatmap analysis of the expression of genes related to inflammation and cytokines/chemokines. (G) Heatmap analysis of DEG in the members of MAPK family. (H) The protein levels of activated JNK (p-p54 and p-p46) and p38 in HUVECs after treatment with different concentrations of Neratinib; n=6 biological independent repeats for JNK and n=5 for p38. (I) Immunofluorescence of JNK and p-JNK in the aortic sinus. Scare bar: 100 μm, n=10 for each group in male *Ldlr*^−/−^ mice and n=7 for each group in female *Ldlr*^−/−^ mcie. (J) Western blot results of p-ASK1 and ASK1 in Neratinib-treated HUVECs exposed to TNF-α; n=6 biological independent repeats for each group. (K) Immunofluorescence of p-ASK1 in HUVECs after treatment with 5 μM Neratinib and stimulation with TNF-α; n=5 biological independent repeats for each group. Sclar bar: 25 μm. (L and M) CETSA experiment; n=3 biological independent samples for L and n=4 for M. (N) DARTS experiment; n=12 biological independent samples for each group. (O) The result of molecular docking of Neratinib and ASK1. (P) The structure of ASK1 includes an N-terminal domain, an ATP binding domain and a kinase activity domain. The N-terminal domain mediates the dimerization of ASK1 under stimulation by TNF-α or ROS. (Q) Dimerization of ASK1 after treatment with Neratinib; n=3 biological independent samples for each group. (Q) Surface plasmon resonance (SPR) analysis. Abbreviations: CETSA: Cellular thermal shift assay, DARTS: drug affinity responsive target stability, Ner: Neratinib, SPR: surface plasmon resonance. All data are presented as the means ± SEMs. Data of p-JNK^+^ area in male mice in I are analyzed using unpaired Student’s t-test with Welch’s correction. Data of p-JNK^+^ area in female mice in I are analyzed using unpaired Student’s t-test. Data of L and N are analysed using paired Student’s t-test. Data of M are analyzed using one-way ANOVA test.

### ASK1 as the molecular target of Neratinib

To identify the molecular targets of Neratinib, we performed transcriptome profiling in Neratinib-treated HUVECs in the presence of TNF-α (Figure. 4A and 4B). In endothelial cells treated with Neratinib, the expression of 992 genes, identified using the criteria of *q* value < 0.05 and log_2_(FC) < −1, was significantly downregulated (Figure. 4C) while 665 genes were significantly upregulated. KEGG enrichment pathways analysis of down-regulated genes related to endothelial dysfunction and atherosclerosis are MAPK signalling pathway, cytokine-cytokine receptor interaction, TNF-α signalling pathway, chemokine signalling pathway and cell adhesion molecules, among which MAPK pathway is the driving common mechanism for the rest of enriched pathways (Figure. 4D). In detail, the transcription levels of adhesion molecules and chemokines were significantly decreased by Neratinib treatment (Figure. 4E and 4F). The transcription of molecules in the MAPK pathway was also significantly decreased after Neratinib treatment (Figure. 4G). These data implied that Neratinib may target MAPK pathway to inhibit endothelial inflammation. Further research in cultured endothelial cells revealed that JNK and p38 phosphorylation was strongly inhibited after Neratinib treatment (Figure. 4H). Immunofluorescence staining of mouse aortic lesions revealed that JNK activation was inhibited in the Neratinib-treated mice (Figure. 4I). Therefore, we next examined whether the activation of ASK1, a protein upstream of JNK and p38^23^, underlies Neratinib’s anti-inflammatory effects. Expectedly, the results revealed that ASK1 activation was significantly reduced in endothelial cells after TNF-α treatment (Figure. 4J and 4K). These findings suggest that Neratinib may reduce endothelial inflammation by inhibiting the ASK1/JNK/p38 pathway. To approach whether Neratinib exhibits anti-inflammatory effects by direct interaction with ASK1, cell thermal shift assay (CETSA) and drug affinity responsive target stability (DARTS), two methods used for assessment of drug‒protein binding, demonstrated that ASK1 protein becomes more stable after incubation with Neratinib (Figure. 4L-N). The domain structure of ASK1 was shown in Figure 4O, including N-terminal domain, ATP binding domain and Kinase activity domain. To investigate the potential binding site(s), we used molecular docking to exhibit that the combination score was −6.48 kcal/mol and that D756, K708 and V757 were pivotal amino acid residues for the binding of Neratinib to ASK1 (Figure. 4P). Further, we observed a Kd value of 2.12 × 10^−6^ M between Neratinib and ASK1 fragment (681-936 aa) binding assessed by surface plasmon resonance (SPR) (Figure. 4P). These data revealed the direct interaction between Neratinib and ASK1 via its ATP binding domain.

In further experiments, we investigated the role of the ASK1/JNK/p38 pathway in Neratinib-mediated protective effects against endothelial cell inflammatory response. We constructed an adenovirus carrying human *MAP3K5* (which encodes *ASK1*). The results demonstrated that the inhibitory effects of Neratinib on *VCAM1* and *SELE* gene expression were partially abrogated (Figure. 5A). The inhibitory effects of Neratinib on VCAM1 protein level was also abrogated (Figure. 5B). We also observed that ASK1 overexpression partially reversed Neratinib-mediated inhibition of monocyte adhesion to endothelial cells (Figure. 5C). In addition, we observed that Selonsertib and GS-444217 (two ASK1 inhibitors) significantly inhibited the upregulation of VCAM1 in HUVECs (Figure. 5E). GS-444217 reduced THP-1 adhesion to HUVECs in a concentration-dependent manner (Figure. 5F and 5G). These results demonstrated that Neratinib mitigated endothelial cell inflammatory response by direct binding to ASK1 and inhibited ASK1 activation and ensuing inflammatory response.

**Figure 5.**
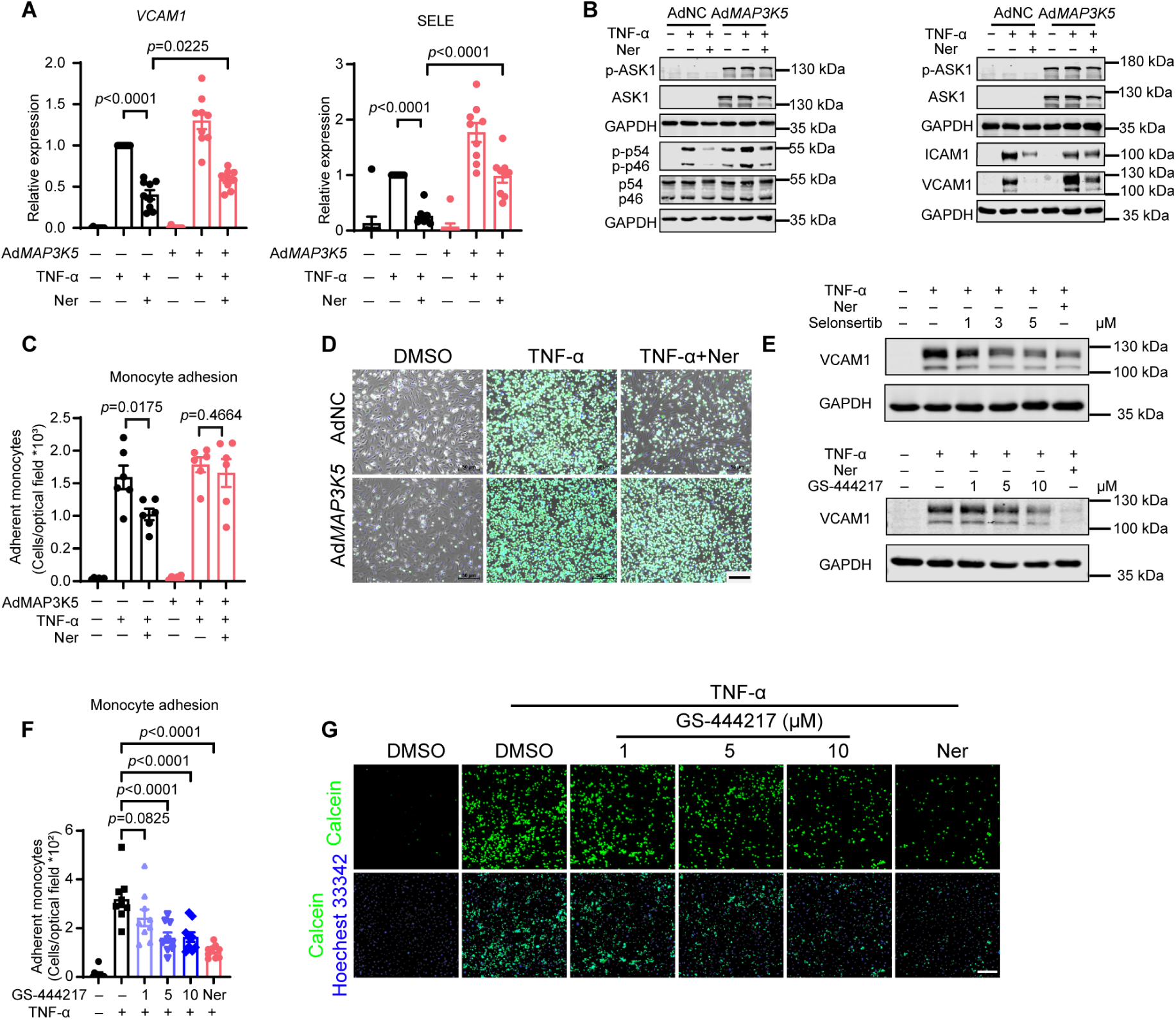
Neratinib reduces the endothelial cell inflammatory response by inhibiting ASK1. (A) The expression of V*CAM1* and *SELE* in HUVECs treated with 5 μM Neratinib in the presence of control or *MAPK3K5* (*ASK1*)-overexpressing adenovirus; n=9 biological independent repeats. (B) Western blot analysis of indicated protein expression in cells treated as described in A; n=3 biological independent repeats. (C and D) Monocyte adhesion assay; n=6 biological independent samples. Scare bar: 50 μm. Adherent THP-1 cells are stained with Calcein-AM. (E) Western blot analysis of VCAM1 in HUVECs treated with escalating doses of two independent ASK1 inhibitors Selonsertib and GS-444217, n=3 per each treatment. (F and G) Monocyte adhesion assay of HUVECs treated with GS-444217, n=9 biological independent repeat samples. All data are presented as the means ± SEMs. Data of A and F are analyzed using one-way ANOVA test. Data of C are analysed using Brown-Forsythe and Welch ANOVA test.

### Rosuvastatin-Neratinib combination therapy provides synergistic anti-atherosclerotic effects

On the basis of the above results, daily administration of 20 mg/kg Neratinib significantly reduced HCD-induced atherosclerosis in *Ldlr^−/−^* mice by inhibiting vascular inflammation without affecting lipid profile. This prompts us to evaluate whether combined Rosuvastatin with Neratinib therapy can prevent atherosclerosis in *Ldlr*^−/−^ mice in a synergistic manner (Figure. 6A and 6B). The results showed that Neratinib and rosuvastatin monotherapy reduced plaque area and lipid deposition (Figure. 6C and 6D). Interestingly, when Neratinib and rosuvastatin were used in combination, there was a significant reduction in plaque area and lipid deposition as compared to Rosuvastatin monotherapy. HE staining and Masson staining revealed that the size of the necrotic core in the aortic sinus plaque was significantly reduced (Figure. 6E), but the collagen content was not significantly increased (Supplement Figure. 5E). Immunofluorescence staining of plaques revealed a sharp decrease in CD68-positive areas after combination therapy (Figure. 6F). To better understand whether neratinib add-on therapy renders a decreased inflammatory profile in Rosuvastatin-treated mice. We performed an Olink proteomics analysis of mouse serum. The results shows that the levels of multiple cytokines (CCL2, CCL4, CSF2, CXCL9, IL7 and IL16) were reduced by Neratinib treatment (Figure. 6G and 6H). These findings indicate that the combination of Rosuvastatin and Neratinib harnesses the advantages of both drugs to synergistically reduce atherosclerosis via a complementary mechanism.

**Figure 6.**
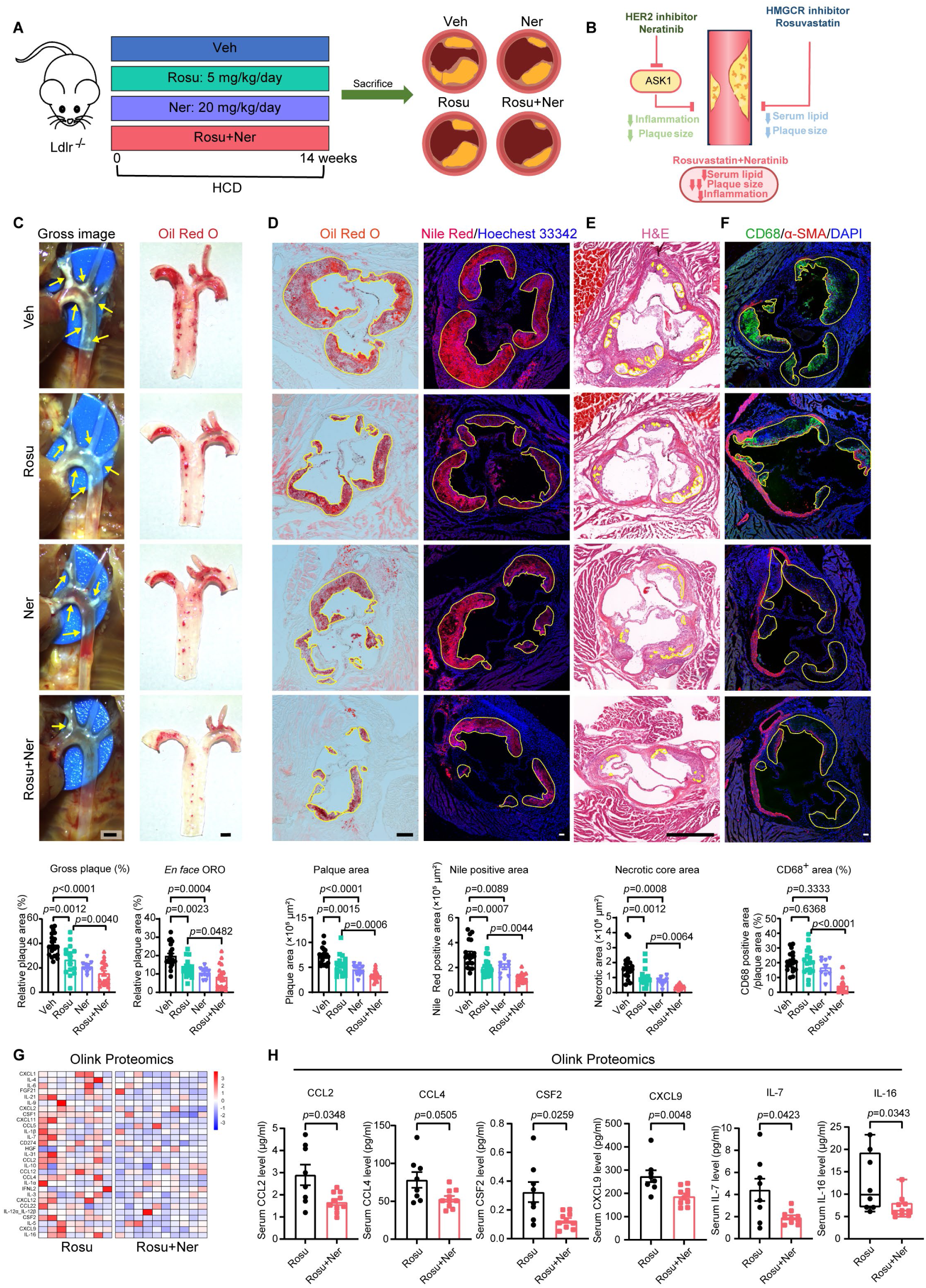
Rosuvastatin and Neratinib combination therapy produced synergistic anti-atherosclerotic effects. (A) Scheme of study design. (B) Graphic of the benefit of combination therapy (C) Gross image and Oil Red O-stained image of *en face* aorta. Scale bar: 1 mm, n=10 for Neratinib group, n=17 for Rosuvastatin, and n=19 for vehicle and combination group in gross image and n=10 for Neratinib group, n=18 for Rosuvastatin, and n=20 for vehicle and combination group in *en face* ORO. (D) Oil Red O staining of aortic sinus. Scale bar: 200 μm, n=10 for Neratinib group, n=18 for Rosuvastatin, and n=20 for vehicle and n=19 for combination group. Nile red staining of neutral lipids in aortic sinus. Scale bar: 50 μm, n=8 for Neratinib group, n=17 for Rosuvastatin, n=20 for vehicle and n=19 for combination group. (E) H&E staining of aortic sinus to indicate the necrotic core area. Scale bar: 625 μm, n=10 for Neratinib group, n=18 for Rosuvastatin, and n=20 for vehicle combination group. (F) Immunofluorescence staining of CD68 and α-SMA in the aortic sinus. Scale bar: 50 μm, n=10 for Neratinib group, n=119 for Rosuvastatin, n=20 for vehicle and combination group. (G and H) Olink proteomics illustrating differentially secreted inflammation associated cytokines in the combination group vs rosuvastatin group, n=8 for Rosuvastatin and n=10 combination group. Abbreviations: ROSU: Rosuvastatin, Ner: Neratinib. All data are presented as the means ± SEMs. Data of C, ORO in D, E and F are analysed using Kruskal-Wallis test. Data of Nile Red staining in D are analyzed using Brown-Forsythe and Welch ANOVA test. Data of H were analysed using unpaired Student’s t-test with Welch’s correction in IL-7, CCL2, CCL4, and CSF2, and unpaired Student’s t-test in CXCL19. Data of IL-16 in H were analyzed using Mann-Whitney test.

## DISCUSSION

Atherosclerosis is a chronic disease driven by dyslipidemia and inflammation. Thus, discovery of inflammation-targeted drugs will complement lipid-lowering based standard care to address the residual inflammatory risk. To this end, drug repositioning is an effective approach to solve the dilemma of high investment and low success rate of new drug development^7^. CMap is a database of gene expression changes caused by extensive numbers of drugs in a variety of human cells^9, 24^. CMap constructs the bridge between drugs, genes and diseases^10^, which could assist to identify new indications of FDA approved drugs. In this study, we discovered a novel anti-inflammatory drug by combined use of in silico and experimental approaches. Neratinib reduces endothelial inflammation via ASK1 inhibition, rather than its classical target HER2/ERBB2. The anti-inflammatory effects of Neratinib translate into atheroprotective effects in both male and female *Ldlr^−/−^* mice. More importantly, add-on therapy of Neratinib to lipid-lowering rosuvastatin conferred synergistic anti-atherosclerotic effects. Therefore, the present study exemplified the use of CMap screening to identify repurposed drug in cardiovascular therapeutics and suggests that Neratinib could potentially serve as a repurposed drug for the treatment of atherosclerosis.

Neratinib is an orally bioavailable, small-molecule tyrosine kinase inhibitor that irreversibly inhibits pan-HER tyrosine kinases (EGFR, HER2/ERBB2, and HER4/ERBB4). Neratinib (240 mg daily), was approved by the FDA for the treatment of HER2-positive breast cancer^25, 26^. Previous studies have focused on the interaction between Neratinib and tumor cells. For example, Neratinib inhibits the growth of breast cancer cells and pancreatic cancer cells^27, 28^. Previous studies have shown that Neratinib inhibits angiogenesis in chick embryos^29^, indicating the possibility of Neratinib as a new antiangiogenic agent. Neratinib also inhibited the activation of hepatic stellate cells and liver fibrogenesis in a CCl_4_-induced mouse model by interrupting the FGF2 signaling pathway^30^. However, the effect and mechanism of action of Neratinib in endothelial cell function and atherosclerosis has not been reported. In this study, we selected 20 mg/kg/d as the dose to treat *Ldlr*^−/−^ mice. The selection criteria were based on: (1) dosage conversion between human and mice^31^; (2) Neratinib at 100 mg/kd/d causes gut injury^32^. Since some tyrosine kinase inhibitors have been reported to cause cardiovascular toxicity, we also assessed the safety of Neratinib in normal C57BL/6J mice and hypercholesterolemia mice. Our data suggested that 20 mg/kg/d can be safely tolerated in mice under normal and diseased conditions evidenced by no significant histological changes and serum biochemistry parameters of major tissues or organs. We then embarked on the assessment of pharmacological actions of Neratinib in vitro and in vivo. We observed that the anti-inflammatory effects of Neratinib exist in different inflammatory conditions as well as in both venous and artery endothelial cells. Finally, the anti-inflammatory effects translate into lower macrophage infiltration in the plaques. Together, these evidences underscore the clinical translational potential of Neratinib.

Neratinib is pan-HER tyrosine kinases inhibitor, however, HER family members expression in endothelial cells can barely be detected despite HER2 expression is relatively enriched in endothelial cells. Intriguingly, genetic intervention (gain or loss-of-function) and pharmacological inhibition of HER2 by other inhibitors did not yield any anti-inflammatory effect, nor abrogating Neratinib’s anti-inflammatory effects, suggesting that HER2 is dispensable for the anti-inflammatory effects of Neratinib. Based on our transcriptomic profiling data, we observed the enrichment of genes in the MAPK pathway. Previous literature has demonstrated that ASK1 represents an important target in inflammatory response and vascular disorders. It has been well established that ASK1 is a MAPK kinase functioning upstream of JNK and p38^23^. The ASK1/JNK/p38 signaling pathway is closely related to thrombosis^33^, vascular inflammation^34–36^ and a variety of metabolic diseases. For example, the ASK1/JNK/p38 signalling pathway is involved in the regulation of SR-A expression in macrophages, which is activated by oxLDL and mediates the formation of macrophage foam cells^37, 38^. However, inhibition of the ASK1/JNK pathway reduces MMP9 production in macrophages and promotes plaque stability^39^. Further, ASK1 mediates smooth muscle cell hypertrophy^40^ and cell survival under stimulation by TNF-α and vascular injury^34, 41^. ASK1 plays an important role in neointimal formation^42^. Loss of ASK1 attenuates vascular injury-induced neointimal hyperplasia^41^. In endothelial cells, ASK1 mediates oxLDL-induced ER stress and endothelial cell injury. The inhibition of ASK1 by GS-4997 alleviated ox-LDL-induced ROS generation and inflammation in endothelial cells^43^. In addition, the inhibition of ASK1 reduces fatty liver and liver fibrosis^44^ and alleviates cardiac hypertrophy^45^. These evidences suggest that ASK1 represents a potential therapeutic target in a plethora of pan-vascular and pan-metabolic diseases. In this study, we characterized the molecular target of Neratinib by SPR and confirmed the direct binding of Neratinib to ASK1. Further cellular experiments revealed that Neratinib attenuates endothelial inflammation by inhibiting ASK1activation, independent of classical HER2 inhibition. In addition, JNK and p38 MAPK activation has been previously shown to be activated following inflammatory stimulation via ASK1, further supporting our hypothesis that Neratinib antagonizes the ASK1/JNK and p38 MAPK signaling cascade to exert its anti-inflammatory effects. Since the dimerization of ASK1 is well known to drive inflammation, detailed mapping of key interacting residues essential for Neratinib to inhibit ASK1 activation and dimerization are warranted. Additionally, considering that breast cancer is related to a higher risk of CVD mortality^46^ and that the cardiovascular toxicity of antineoplastic drugs^47^, such as pertuzumab^48^, trastuzumab^49^ and bevacizumab^50^, the protection of Neratinib against atherosclerosis provides a potentially safer drug recommendations for patients with breast cancer.

By using a variety of molecular and cellular approaches as well as in vivo animal models, we demonstrate that Neratinib attenuates endothelial inflammation and atherosclerosis and ASK1 inhibition mediates the anti-inflammatory effects of Neratinib. However, we have to recognize there are several limitations in this study. First, we cannot exclude the possibility that other signaling pathways may also contribute to the anti-inflammatory effects of this drug. Second, whether HER2 inhibition contribute to Neratinib-mediated atheroprotective effects in vivo, in particular in female *Ldlr*^−/−^ mouse bearing breast cancers has not been assessed. Last but not least, we discovered that Neratinib protect against atherosclerosis in *Ldlr*^−/−^ mouse. Considering the difference between mouse and human, whether the protective effects of Neratinib in mice can be translatable to humans is unknown.

In summary, the present study identifies Neratinib as a novel repurposed drug against endothelial inflammation and atherosclerosis. Future clinical studies are warranted to determine the therapeutic actions, tolerability and safety of Neratinib in patients with atherosclerosis

## MATERIALS AND METHODS

### Animal and treatment

B6/JGpt-*Ldlr*^em1Cd82^/Gpt (*Ldlr*^−/−^) mice were purchased from GemPharmatech (T001464; Jiangsu, China) and housed in isolated ventilated cages (IVCs) in a temperature- and humidity-controlled animal facility with a 12/12-hour light/dark cycle at 23°C. The mice had free access to water and food. All animal experiments were approved by the Animal Research Ethics Committees of the University of Science and Technology of China (animal ethics no. USTCACUC212401038. Animal ethics application is accepted at 2021-02-02.). The animals were managed and used in accordance with the National Institutes of Health Guide for the Care and Use of Laboratory Animals.

### *Ldlr*^−/−^ mice at 8 weeks of age were randomly divided into indicated groups

Atherosclerosis was established by feeding them with a high-cholesterol diet (HCD, D12108C; Research Diet, New Brunswick, NJ, USA) for 14 weeks. Neratinib was purchased from TargetMol (T2325; Boston, Massachusetts, USA) and was administered orally at a dosage of 20 mg/kg/day. The vehicle used was 0.5% carboxymethylcellulose sodium (C9481; Sigma‒Aldrich; Missouri, USA) +0.4% Tween-80 (A600562; Sangon Biotech; Shanghai, China). Sex as a biological variable was considered in the study design. Both male and female *Ldlr*^−/−^ mice were used for assessing preventive effects of Neratinib. In the combination experiment, the same dose of Neratinib was used for the mice, which were concurrently orally administered 5 mg/kg/day Rosuvastatin (MedChemExpress, HY-17504A, Shanghai, China). All the mice were dosed daily and sacrificed after 14 weeks of HCD feeding to assess atherosclerotic plaque development and composition. *En face* aorta were fixed with 4% paraformaldehyde and stained with Oil Red O to visualize atherosclerotic plaques. Mouse livers and hearts were fixed with 4% paraformaldehyde (P0099; Beyotime, Shanghai, China) and embedded in OCT compound to obtain tissue sections before they were processed for Oil Red O and H&E staining. Blood was collected retro-orbitally and centrifuged at 12,000 × g for 10 minutes at 4°C to obtain serum for analysis of blood biochemistry parameters or Olink proteomics assay. Serum and other tissues were stored at −80°C.

### Statistical analysis

All the data were analysed via GraphPad 9 software (GraphPad, San Diego, USA) and are presented as the means ± SEM. All the in vitro experiments were repeated at least three times. N number representing biological replicates was specified in each figure legend. The *p* values for all the data were calculated via an unpaired two-sided Student’s t test or one-way ANOVA. For all the data, *p*< 0.05 indicated statistical significance.

Detailed materials and methods are available in online data suppplement.

## List of Supplementary Materials

Materials and Methods

Fig S1 to S5

Tables S1 to S9

References (*51-58*) (numbers for references only cited in SM)

## Acknowledgments

The authors would like to thank Shuo Chen, Zhenghong Liu, Zhidan Zhang, Meijie Chen, Yaping Zhao, Chenyang He, Monan Liu, and Iqra Ilyas for assisting tissue harvest. S.X. is Senior Humboldt Research Fellow of Alexander von Humboldt Foundation, Germany. This study was supported by grants from the National Key R&D Program of China (Grant No. 2021YFC2500500), the National Natural Science Foundation of China (Grant Nos. 82370444, 82070464, 12411530127). This work was also supported by the Program for Innovative Research Team of The First Affiliated Hospital of USTC (CXGG02) and Anhui Provincial Natural Science Foundation (Grant No. 2208085J08).

## Data and materials availability

All data generated and used in this study are either included in this article (and its Supplementary Information) or are available from the corresponding author on reasonable request. Data will be made available on request. RAW data of RNA-seq was deposited in Genome Sequence Archive (Genomics, Proteomics & Bioinformatics 2021) in National Genomics Data Center, China National Center for Bioinformation / Beijing Institute of Genomics, Chinese Academy of Sciences (GSA: CRA016475) that are publicly accessible at https://ngdc.cncb.ac.cn/gsa. The public data we used in Supplement Figure. 4A and 4B are from Gene Expression Omnibus with accession number: PRJNA646233 and GSE159677.

## Author contributions

Conceptualization: Suowen Xu, Jianping Weng, Fanshun Zhang

Methodology: Fanshun Zhang, Yanjun Yin, Zhihua Wang, Xiumei Wu

Investigation: Fanshun Zhang

Data analysis: Fanshun Zhang, Zhihua Wang

Funding acquisition: Suowen Xu, Jiangping Weng

Project administration: Suowen Xu, Jiangping Weng

Supervision: Suowen Xu

Writing – original draft: Fanshun Zhang

Writing – review & editing: Suowen Xu, Danielle Kamato

## Supplementary Materials for

**The PDF file includes:**

Supplementary Materials and Methods

Figs. S1 to S5

Tables S1 to S9

References (51-58)

## Materials and Methods

### Animals and treatment

B6/JGpt-*Ldlr*^em1Cd82^/Gpt (*Ldlr*^−/−^) mice were purchased from GemPharmatech (T001464; Jiangsu, China) and housed in isolated ventilated cages (IVCs) in a temperature- and humidity-controlled animal facility with a 12/12-hour light/dark cycle at 23°C. The mice had free access to water and food. All animal experiments were approved by the Animal Research Ethics Committees of the University of Science and Technology of China (animal ethics no. USTCACUC212401038). The animals were managed and used in accordance with the National Institutes of Health Guide for the Care and Use of Laboratory Animals.

For the toxicology experiment, male mice at 8 weeks on the background of C57BL/6J mice were randomly divided into three groups, n=5 for each group. Neratinib was purchased from TargetMol (T2325; Boston, Massachusetts, USA) and was administered orally at a dosage of 20 mg/kg/day and 60 mg/kg/day. The vehicle used was 0.5% carboxymethylcellulose sodium (C9481; Sigma‒Aldrich; Missouri, USA) +0.4% Tween-80 (A600562; Sangon Biotech; Shanghai, China). After 4 weeks, mice were sacrificed to estimate safety. We measure organic weight and detect serum biochemistry parameters related to heart, liver and kidney.

For the atherosclerosis treatment experiment, *Ldlr*^−/−^ mice at 8 weeks of age were randomly divided into two groups, and atherosclerosis was established by feeding them a high-fat high-cholesterol diet (HCD, D12108C; Research Diet, New Brunswick, NJ, USA) for 14 weeks. Neratinib was administered orally at a dosage of 20 mg/kg/day. The vehicle used was 0.5% carboxymethylcellulose sodium +0.4% Tween-80. In the combination experiment, the same dose of Neratinib was used for the mice, which were orally administered 5 mg/kg/day Rosuvastatin. Rosuvastatin was purchased from MedChemExpress (HY-17504A; Shanghai, China). All the mice were sacrificed after 14 weeks to analyse the process of atherosclerosis. Aortic arteries were fixed with 4% paraformaldehyde and stained with Oil Red O to visualize atherosclerotic plaques. Mouse livers and hearts were fixed with 4% paraformaldehyde (P0099; Beyotime, Shanghai, China) and embedded in OCT compound to obtain tissue sections before they were processed for Oil Red O and H&E staining. Blood was collected via eyeball extraction and centrifuged at 12,000 × g for 10 minutes at 4°C to obtain serum for analysis of blood biochemistry parameters. Serum and other tissues were stored at −80°C.

### Determination of blood biochemistry parameters

Serum was collected for analysis of levels of serum lipids and inflammatory cytokines (by Olink Proteomics described as below) and for evaluating the organ toxicity of drugs. Total cholesterol (CHO), total triglyceride (TG), high-density lipoprotein (HDL) and low-density lipoprotein (LDL) levels were determined colorimetrically as index of lipid profile. Alanine transaminase (ALT), aspartate transaminase (AST), urea, creatine (CREA), creatine kinase (CK) and lactate dehydrogenase (LDH) were determined for index of liver, kidney and heart function. All tested items were detected by Servicebio via commercially available kits (S03030 for ALT, S03040 for AST, S03036 for UREA, S03076 for CREA, S03024 for CK, S03034 for LDH, S03027 for TC, S03042 for CHO, S03025 for HDL, and S03029 for LDH; Rayto; Guangzhou, China).

### Oil Red O staining

Oil Red O staining of *en face* aorta and aortic sinus cryosections was performed as previously described^51^. Aorta or cryosections of aortic sinus were rinsed with 60% isopropanol (A600918‒0500; Sangon Biotech) for 10 s and stained with 60% Oil Red O (O0625; Sigma‒Aldrich;) for 5 min. Then, tissue samples were rinsed with 60% isopropanol for 5 min. Finally, the tissues samples were washed with PBS and photographed for analysis of the aortic plaque area. Cryosections were washed with PBS, mounted in glycerol jelly, covered with coverslips and photographed for analysis of the sinus plaque area using Image J software.

### Histological staining and Masson staining

Mouse livers were fixed with 4% paraformaldehyde overnight and embedded in paraffin (8 μm thick sections were cut with a Leica RM2235). Histological staining and Masson staining of the sections were entrusted to Ribiogy by using H&E staining commercially available kits (6765001, 6766010, Thermo Scientific; Massachusetts, USA) and Masson staining kits (C0189S; Byotime). The images were obtained with a Pannoramic 250/MIDI to analyze the necrotic core size and collagen-positive area.

### Cell culture

Primary human umbilical vein endothelial cells (HUVECs) and human artery endothelial cells (HAECs) were purchased from LifeLine Cell Technology (CA, USA) (FC-003 for HUVECs; FC-0014 for HAECs). Cells were grown in endothelial cell culture medium (ECM, LL-0003; LifeLine). All experiments involving HUVECs and HEACs were performed with passage 4–6. HUVECs and HAECs were treated with 10 ng/ml TNF-α (AF-300-01A; Peprotech; NJ; USA), IL-1β (200-01B; Peprotech) or 1 μg/ml lipopolysaccharide (LPS, L2630; Sigma‒Aldrich) to establish cellular models of endothelial inflammation when the cells reached 90% confluence. Human monocyte cells (THP-1) were purchased from American Type Culture Collection (ATCC) (TIB-202) and cultured in RPMI-1640 medium (KEL1501-500; KeyGen BioTECH, Jiangsu, China) supplemented with 10% FBS (12103C; Millipore Sigma). The cells were cultured in 10 mm dishes, seeded in 12-well plates and incubated in a 5% CO_2_ atmosphere at 37°C for experiments.

### Monocyte adhesion assay

HUVECs or HAECs were passaged into 12-well plates, treated with Neratinib and stimulated with TNF-α. THP-1 cells were suspended in 2 ml of RPMI-1640 and stained with Calcein AM (C2012; Beyotime) for 30 min at 37°C. The THP-1 cells were then washed with RPMI-1640, and the HUVECs were washed with PBS three times. The THP-1 cells were added to the plates and incubated with HUVECs for 30 min. Finally, the plates were washed with PBS three times to get rid of the unadhered THP-1 cells. Three different optical fields were recorded randomly via a microscope (Axiovert 5/Axiovert 5 digital; ZEISS, Germany). The adherent cells were counted via ImageJ software (Bethesda, USA).

### Connective Map (CMap) screening

CMap database was designed and carried out according to established procedures^52^. Briefly, the process of CMap exploration is described. RNA-sequencing transcriptomic data of primary human umbilical vein endothelial cells treated with TNF-α or IL-1β were analyzed to select top 150 upregulated and downregulated genes with the criteria of *q* value < 0.05 and log_2_(FC) > 1 or log_2_(FC) < −1 (Supplement Table 1 and Supplement Table 2). The top 150 genes were constructed as the inflammation dataset, which were uploaded to the CMap database, matched and scored to obtain drug perturbed signature in inverse relation to that of TNF-α and IL-1β. We defined the parameter “S” to represent the score of the compound in lowering TNF-α and IL-1β induced gene signature. We selected 24 compounds based on the value of “S” for verification of anti-inflammatory effects in cultured endothelial cells (Supplement Table 3). Under ideal circumstances, the top effective anti-inflammatory compound has higher score in each set. In addition, the scores between different sets are similar.

Cellular screening is to check the anti-inflammatory effects of the compounds. The average ranks (*R̅*) of all the evaluated items was the criterion. A smaller average rank reflected a stronger anti-inflammatory effect.

“S” is computed as follows:

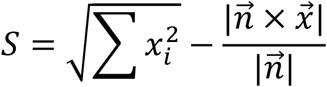

Where *i*, *x*_*i*_, and 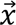 represent the drug number, the score of the drug in the dataset, and the vector consisting of the CMap score, respectively. Where 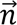 is vector of which each component is one.

The average ranks (*R̅*) is computed as follows:

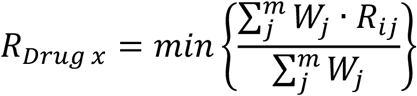

Where *i* and *j* represent the drug number and the item number, respectively. Where *W*_*j*_ and *R*_*ij*_ represent the weight of this item and the rank of the drug in this item. We regard *W*_*j*_ as constant because all the items are in the same level (cellular level). The rank value from small to large reflects the effect of anti-inflammation from strong to weak. The rank values of the drug are listed in Supplement table 4.

### Transcriptome sequencing (RNA-seq)

Total RNA was extracted from HUVECs via an RNA-Quick Purification Kit (RN001; ES Science, Shanghai, China) and stored at −80°C. The absorbance of the RNA was detected with a NanoDrop 2000 (Thermo Fisher; USA) and OD260/OD280 values of RNA was between 2.0∼2.2, after which the samples were sent to Gene Denovo Biotechnology Co. (Guangzhou, China) for sequencing. The mRNA was enriched by magnetic beads with oligo(dT). The first strand of cDNA was synthesized using the fragmented mRNA as a template and random oligonucleotides as primers to construct the library. The resulting library was sequenced via Illumina NovaSeq 6000.

### Accession number

The raw sequencing data of Neratinib-treated HUVEC reported in this paper have been deposited in the Genome Sequence Archive (Genomics, Proteomics & Bioinformatics 2021) in National Genomics Data Center, China National Center for Bioinformation / Beijing Institute of Genomics, Chinese Academy of Sciences (GSA: CRA016475) that are publicly accessible at https://ngdc.cncb.ac.cn/gsa. The public data we used in Supplement Figure. 4A and 4B are from Gene Expression Omnibus with accession number: PRJNA646233 ^53^ and GSE159677 ^54^.

### Olink Target 48-cytokine mouse panel

Olink proteomics was supported by China National BGI (Guangdong, China). The Target 48 cytokine Mouse Panel from Olink (Uppsala, Sweden) was used to quantify the absolute concentrations of 43 proteins for each sample (Sequanta Technologies Co., Ltd). Multiplex proximity extension assay panels were used to quantify each protein. A proximity extension assay uses a dual-recognition immunoassay, in which two matched antibodies labelled with unique DNA oligonucleotides simultaneously bind to the target protein in solution. This brings the two antibodies into proximity, allowing hybridization and serving as a template for a DNA polymerase-dependent extension step. Double-stranded DNA is unique to a specific antigen and is amplified via a primer, which is quantitatively proportional to the plasma concentration of the target protein. The amplified targets were then quantified via RT‒PCR via a Fluidigm BioMarkTM Microfluidic qPCR instrument. The collected data are presented in standard units (pg/mL). Raw data of Olink proteomics are listed in Supplement table 5.

### Surface plasmon resonance for affinity determination

Our prior reports have elaborated on the Surface Plasmon Resonance (SPR) technique utilized in this study^55, 56^. In essence, we leveraged the Biacore T200 platform (GE Healthcare) to quantify the dissociation constants (KD) governing protein-ligand interactions. As a foundational step, the CM5 chip (GE Healthcare) facilitated the immobilization of the target protein. Activation of the chip involved exposure to 1-ethyl-3-(3-dimethylaminopropyl) carbodiimide (EDC, GE Healthcare) and N-hydroxysuccinimide (NHS, GE Healthcare) for 60 seconds at 10 μL/min flow rate, prior to protein binding. Subsequently, ASK1 protein was diluted to approximately 50 mg/mL in a 10 mmol/L sodium acetate buffer (pH 4.5) and introduced into the chip at 10 μL/min for 120 seconds, achieving saturation of the detection channel. Approximately 8000 response units (RU) of ASK1 fragment (681-936 aa) (Ag5818; Proteintech, USA) were coupled per channel. Thereafter, the channel was blocked with ethanolamine at 10 μL/min for 120 seconds to prevent non-specific interactions. To assess the affinity between ASK1 and Neratinib, the compound was sequentially diluted across eight concentrations ranging from 1.25 μM to 0 nM and flowed through the chip, both in the presence and absence of the target protein channel, at 30 μL/min for 180 seconds, with concentrations increasing from lowest to highest. Following each concentration injection, the chip was regenerated using NaOH at 30 μL/min for 60 seconds, facilitating real-time data recording and storage. Ultimately, the Biacore T200 analysis software (GE Healthcare) was employed to analyze and organize the acquired data. Given anomalous responses observed at the two highest concentrations, the KD value was determined by fitting the data from the six lowest concentration points. Raw data of SPR are listed in Supplement table Table 6.

### Real-time quantitative PCR (RT‒qPCR)

Total RNA from tissues and cultured cells was extracted with a Tissue RNA Purification Kit Plus (ES-RN002plus; ES Science), and the absorbance of the RNA was detected with a NanoDrop 2000 spectrophotometer (Thermo Fisher) to obtain values between 2.0∼2.2 and concentrations greater than 50 ng/μl. Next, cDNA was obtained by using a PrimeScript™ RT reagent Kit (RR037Q; Takara Biomedical Technology, Beijing, China). The expression levels of target genes were detected with a LightCycler* 96 Instrument II (Roche, Basel, Switzerland) using TB Green® Premix Ex Taq™ (Tli RNaseH Plus) (RR420A; Takara Biomedical Technology, Beijing, China). The housekeeping gene GAPDH was used for normalization. The fold change relative to the respective controls was calculated via the 2-ΔΔCt method. The primers used in this study were designed by NCBI, synthesized by Sangon Biotechand are listed in Supplemental Table 7.

### Western blot

The protein was extracted with sample loading buffer (20325ES05; Yeasen, Shanghai, China). The protein samples were heated at 95°C for 10 min, separated via 10% SDS‒PAGE and transferred to nitrocellulose blotting membranes (FFN08; Beyotime). The membranes were blocked with Intercept® (PBS) blocking buffer (927-70001; LICOR, USA) for 40 min at room temperature. The membranes were subsequently incubated with the primary antibody overnight at 4°C. The primary antibodies used in this study are shown in Supplemental Table 8. The membranes were incubated with secondary antibody at room temperature after being washed three times with TBS-T (C006161; Sangon Biotech) with 0.1% Tween 20 (A600560; Sangon Biotech, Shanghai, China). The secondary antibodies used were IRDye® 680RD goat anti-mouse IgG (H + L) antibody (925--68070; LICOR) and IRDye® 800RD goat anti-rabbit IgG (H + L) antibody (926--32211; LICOR). After 1 h, the membranes were washed three times with TBS-T. The immunoblot signals were detected via Odyssey® CLx, and densitometrical analysis of the bands were performed with ImageJ software (Bethesda, USA).

### Immunofluorescence assay

The tissue samples were cut into 10 μm sections with a Leica CM1950 cryostat (Leica, Germany). All the sections were placed on glass slides and stored at −80°C. The slides were dried at room temperature for 20 min and washed with PBS to remove the OCT compound. Before they were incubated with the primary antibody, all the slides were blocked with blocking buffer (0.2% Triton X-100, 2% BSA in PBS) for 1 hour at room temperature. The slides were incubated with primary antibodies at 4°C overnight. The slides were washed with PBS three times and incubated with secondary antibodies at room temperature for 1 hour. Finally, the slides were stained with DAPI and covered with Antifade Mounting Medium (P0126; Beyotime). The primary antibodies used were rat anti-mouse CD68 antibody (FA-11; Bio-Rad, USA; 1:500 dilution) and rabbit anti-mouse SMA antibody (14395-1-AP; Proteintech, USA; 1:500 dilution). The secondary antibodies used were goat anti-rat IgG (H+L) antibody (A11006; Invitrogen; 1:1000 ratio dilution) and goat anti-rabbit IgG (H+L) antibody (A11010; Invitrogen; 1:1000 ratio dilution). Fluorescence images were acquired by using Leica TCS SP II (Leica; Germany). The positive areas were analyzed via ImageJ software (Bethesda, USA).

### Small interfering RNA (siRNA), adenovirus and transfection

The siRNAs against *ERBB2 (HER2)* and *MAP3K5 (ASK1)* and the negative control (siNC) were designed from Suzhou Biosyntech (ERBB2 Lot. # ZWJP0103 and MAP3K5 Lot. # ZWJP0801 Jiangsu, China). The sequences of these siRNAs are shown in Supplemental Table 9. An adenovirus encoding human *ERBB2* with a CMV promotor was constructed by WZ Bioscience, Inc. (9.8 × 10^10^pfu/ml; pADM-CMV-FH-FLAG > h*ERBB2*, NM 001005862; Shandong, China). An adenovirus encoding human *MAP3K5* with the EF1A promoter was constructed by VectorBulider (1.44 × 10^10^pfu/ml, pAV-EGFP-EF1A>h*MAP3K5*, NM_005923.4; Guangzhou, China). HUVECs or HEK293T cells were seeded in 12-well plates and transfected siRNA or plasmid with Invitrogen™ Lipofectamine™ 2000 reagent (11668027; Thermo Fisher) following the manufacturer’s instructions. The expression levels of target genes were tested by RT‒qPCR and Western blotting after transfection for 48 hours.

### Cellular thermal shift assay (CETSA) and drug affinity responsive target stability (DARTS)

CETSA^57^ and DARTS^58^ experiment were designed and carried out according to established procedures. For the CETSA experiments, HEK293T cells were transfected with the HA-ASK1 plasmid to overexpress ASK1 for 24 hours and harvested with PBS supplemented with 1x protease inhibitor cocktail. Then, the cell suspensions were freeze-thawed three times with liquid nitrogen and centrifuged to obtain clear supernatants. The cell lysate was incubated with 50 μM Neratinib or DMSO at room temperature for 30 minutes. The cell lysate was divided into 7∼8 aliquots and heated individually at different temperatures for 3 minutes, followed by cooling for 3 minutes at room temperature. After heating, the cell lysates were centrifuged to obtain supernatants, mixed with protein loading buffer, and analysed by SDS‒PAGE followed by western blot analysis.

For DARTS experiments, the cell lysates were obtained in the same way with CETSA experiments, containing 100 μg of protein and incubated with or without 50 μM Neratinib for 30 minutes at room temperature. Pronase from Streptomyces *griseus* (53702; Sigma‒Aldrich, MO, USA) was added to all the samples at a pronase: supernatants protein ratio of 1:1400 for 10 min, and the reaction was stopped with the addition of a 20x protease inhibitor. The lysates were mixed with protein loading buffer and analysed by SDS‒PAGE followed by western blot analysis.

## Supplementary figures

**Supplemental figure 1.**
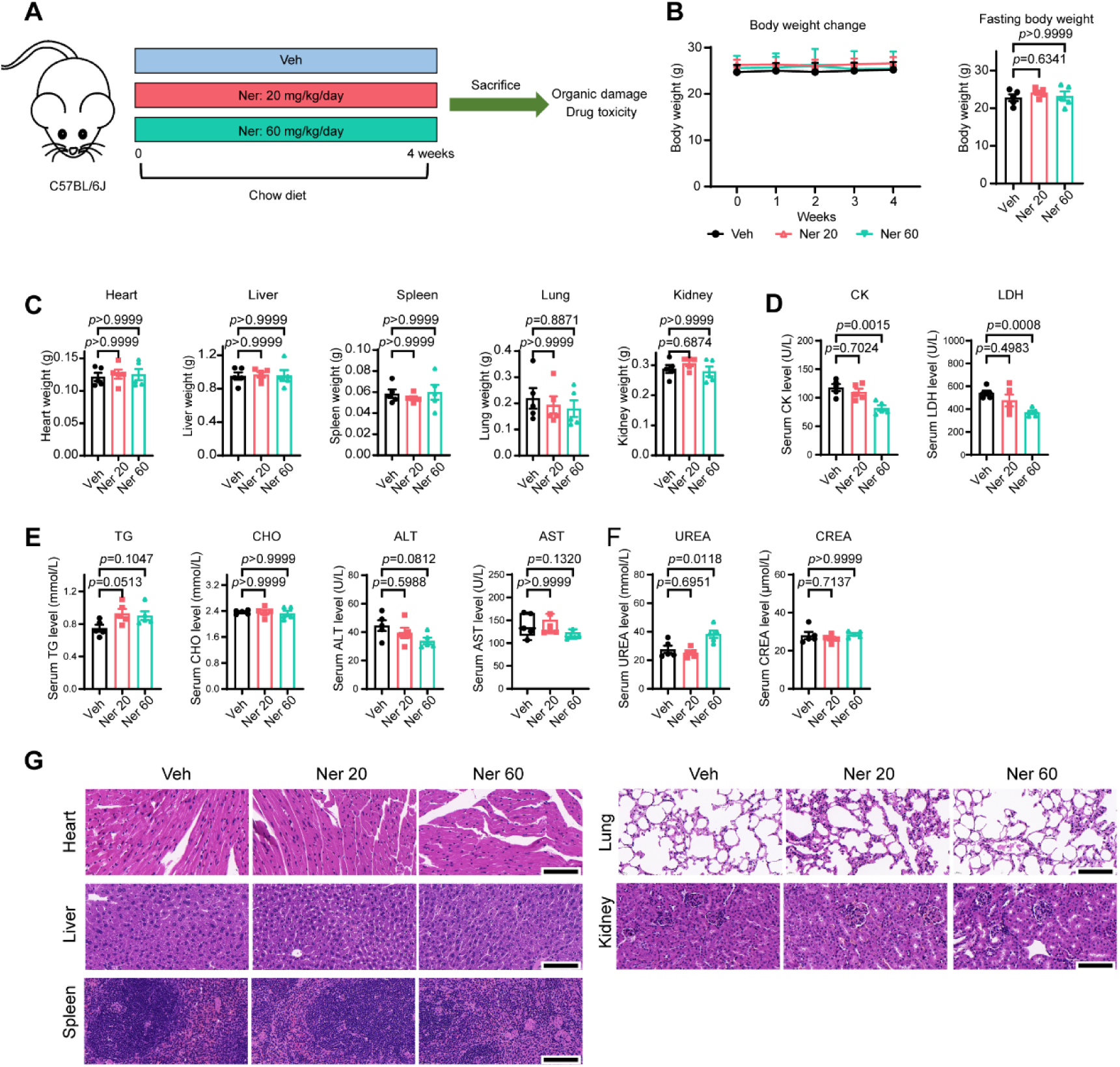
Effect of Neratinib on male C57BL/6J mice fed with normal chow diet. (A) Scheme of study design. Eight-week-old male C57BL/6J mice were divided into three groups and orally administered 0, 20 mg/kg or 60 mg/kg Neratinib every day; n=5 for each group. (B) Body weight curves of the mice from week 1 to 4. (C) Heart, liver, spleen, lung and kidney weight in each group. (D) Serum biochemistry parameters of heart injury-CK and LDH. (E) Serum biochemistry parameters of liver injury-ALT and AST. (F) Serum indices of kidney injury-UREA and CREA. (G) H&E staining of indicated tissues. Abbreviations: ALT: alanine transaminase, AST: aspartate transaminase, CHO: total cholesterol, CK: creatine kinase, CREA: creatine, HDL: high-density lipoprotein, LDH: lactate dehydrogenase, LDL: low-density lipoprotein, Ner: Neratinib, TG: total triglyceride, Veh: vehicle. All Data are presented as the means± SEMs. Data of B, C, TG, CHO, ALT, UREA, CREA and CK in E were analyzed using one-ANOVA test. Data of AST in E were analyzed using Kruskal-Wallis test. Data of LDH in E were analyzed using Brown-Forsythe and Welch ANOVA test.

**Supplemental figure 2.**
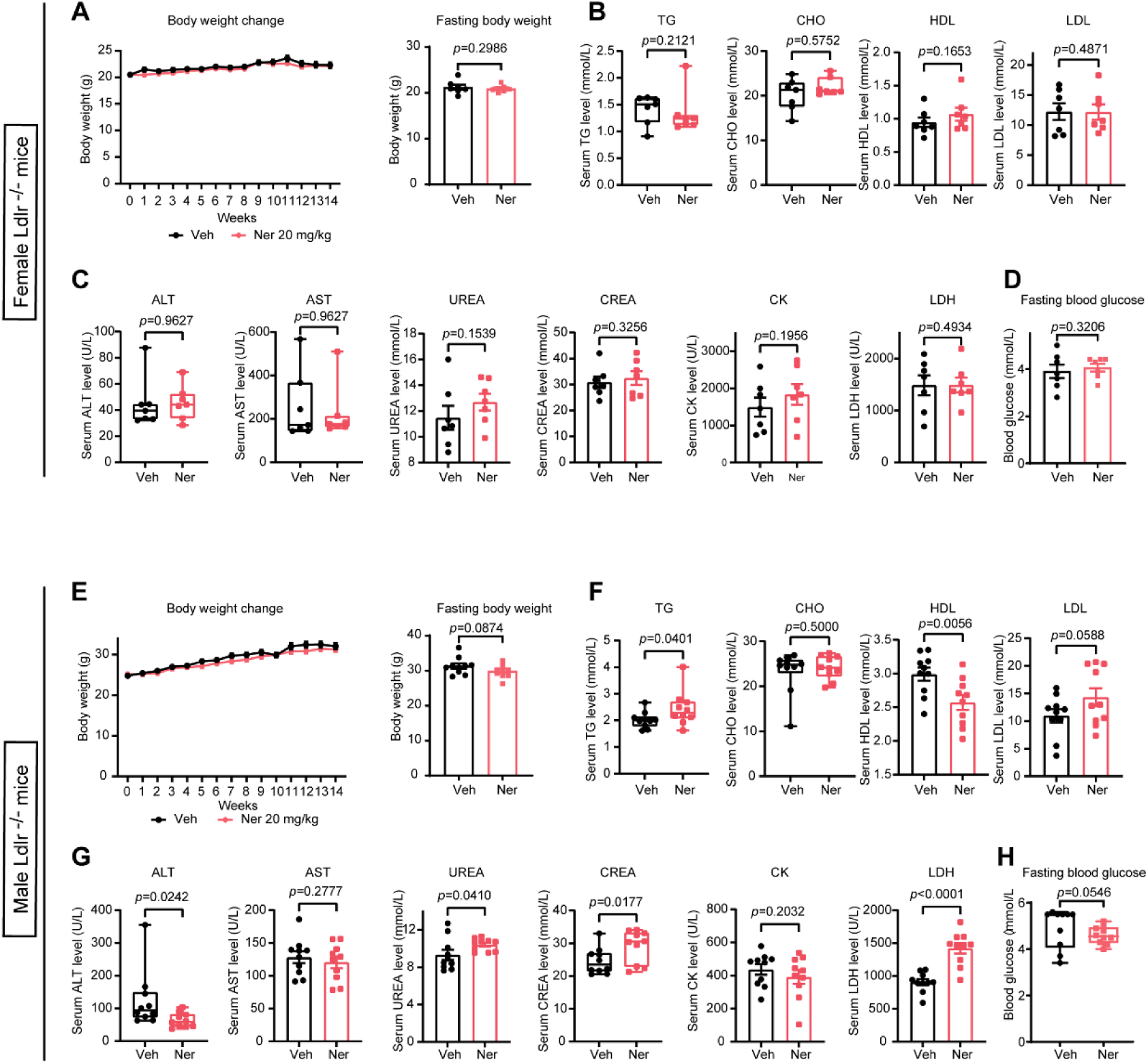
Body weight changes and serum biochemistry parameters of *Ldlr*^−/−^ mice. (A) Body weight curve of female *Ldlr*^−/−^ mice, n=7 for each group in female mice and n=10 for each group in male mice. (B and C) Serum biochemistry parameters of female mice. (D) Fasting blood glucose of female *Ldlr*^−/−^ mice. (E) Body weight curve of male *Ldlr*^−/−^ mice. (F and G) Serum biochemistry parameters of male mice. (H) Fasting blood glucose of male *Ldlr*^−/−^ mice. Abbreviations: ALT: alanine transaminase, AST: aspartate transaminase, CHO: total cholesterol, CK: creatine kinase, CREA: creatine, HDL: high-density lipoprotein, LDH: lactate dehydrogenase, LDL: low-density lipoprotein, Ner: Neratinib, TG: total triglyceride, Veh: vehicle. All data are presented as the means ± SEMs. Data of A, HDL and LDL in B, UREA, CREA, CK and LDH in C, and D are analyzed using unpaired Student’s t-test. Data of TG and CHO in B, ALT and AST in C, and H were analysis using Kolmogorov-Smirnov test. Data of E are analyzed using unpaired Student’s t-test. Data of HDL and LDL in F, AST, CK and LDH in G are analyzed using unpaired Student’s t-test. Data of TG and CHO in F, ALT and CREA in G were analysis using Kolmogorov-Smirnov test. Data of UREA in G were analyzed unpaired Student’s t-test with Welch’s correction.

**Supplemental figure 3.**
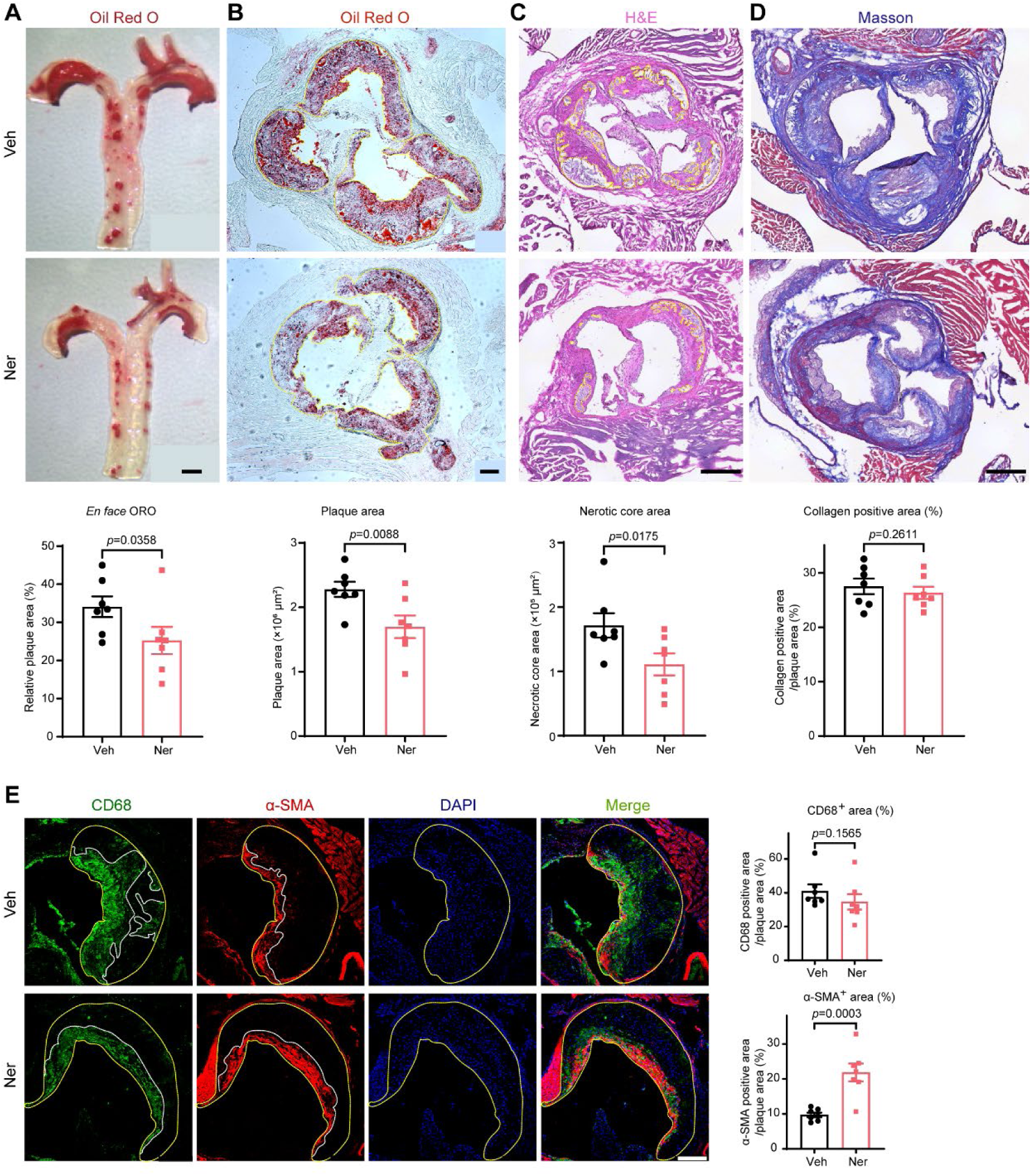
Neratinib prevented atherosclerosis in female *Ldlr*^−/−^ mice. Eight-week-old female *Ldlr*^−/−^ mice were fed a high-fat high-cholesterol diet for 14 weeks and were orally administered with vehicle or 20 mg/kg Neratinib. n=7 in each group. (A) Oil Red O staining of the *en face* aorta. Scale bar: 1 mm. (B) Oil Red O staining of aortic sinus cryosections. Scale bar: 20 μm. (C and D) H&E and Masson staining of aortic sinus sections to indicate the necrotic core area and collagen content. Scale bar:250 μm. (E) Immunofluorescence staining of CD68 and α-SMA in the aortic sinus plaques to indicate infiltrated macrophages and smooth muscle cell content. Scale bar: 200 μm. All data are presented as the means ± SEMs. Data of A-E except α-SMA in E were analyzed with unpaired Student’s t-test. Data of α-SMA in E were analyzed unpaired Student’s t-test with Welch’s correction.

**Supplemental figure 4.**
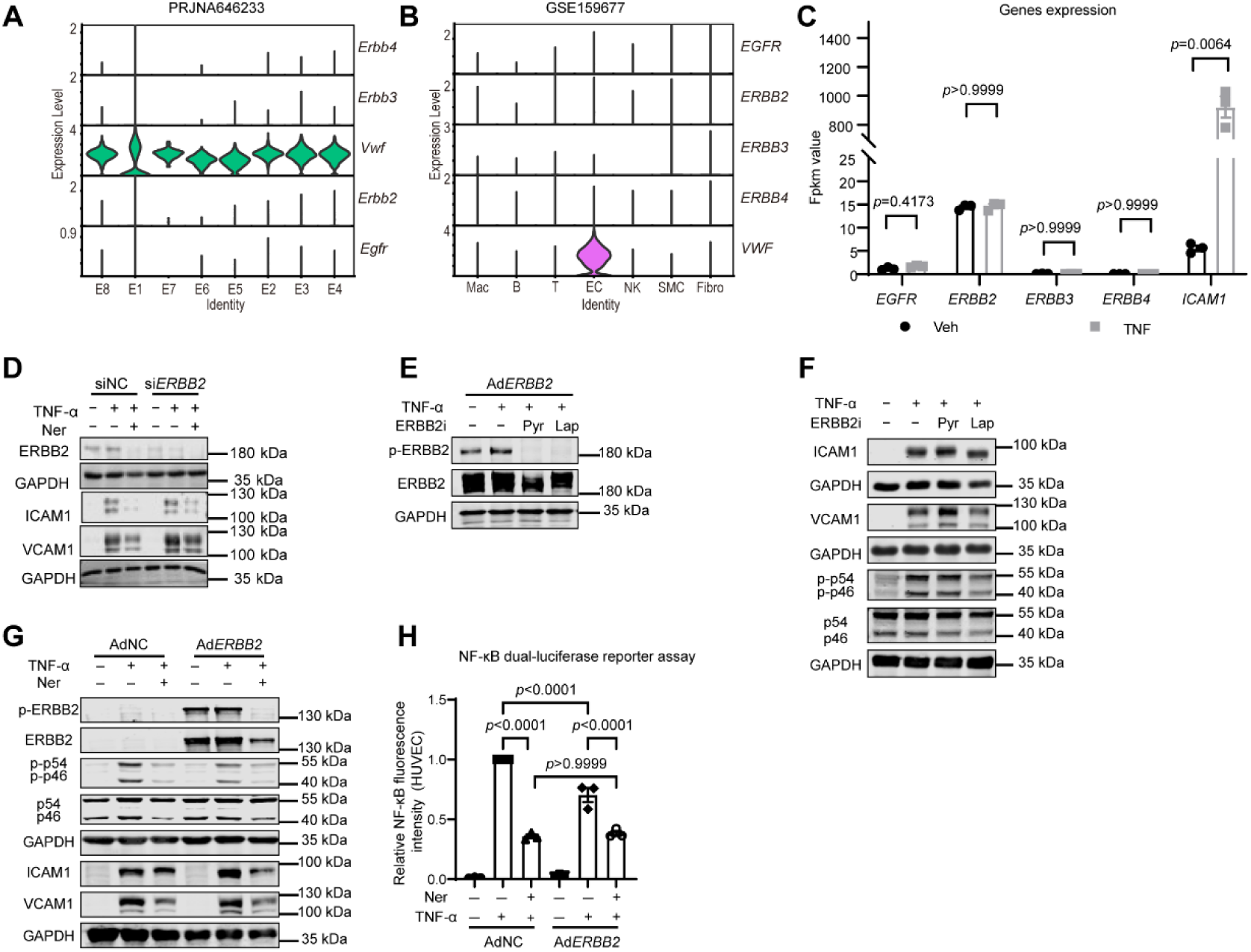
Neratinib reduced endothelial cell inflammatory response independent of the inhibition of HER2/ERBB2. (A) Analysis of single-cell RNA-seq from mouse artery to show the gene expression of *ERBB* family members in different endothelial cell cluster (E1-E8). Data are mined from PRJNA646233 ^53^. (B) Comparative analysis of single-cell RNA-seq from human calcified atherosclerotic core (AC) plaques and patient-matched proximal adjacent (PA) portions of carotid artery to show the gene expression of different *ERBB* family members in endothelial cells and other vascular cell types. Data are mined from GSE159677 ^54^. (C) Gene expression of *ERBB* family members in HUVECs exposed to TNF-α. (D) Effect of *ERBB2* silencing on Neratinib-mediated inhibition of ICAM1 and VCAM1 protein expression; n=3. (E) The protein levels of p-ERBB2 and ERBB2 in HUVECs overexpressed with ERBB2 adenovirus after treatment with 1 μM pyrotinib and 5 μM Lapatinib; n=3 biological independent repeats. (F) The protein levels of ICAM1 and VCAM1 in HUVECs after treatment with two different ERBB2 inhibitors; n=3. (G) The effect of ERBB2 overexpression on Neratinib-mediated inhibition of ICAM1 and VCAM1 protein expression; n=3. Abbreviations: Lap: Lapatinib, Pyr: Pyrotinib. All data are presented as the means ± SEMs. Data of C are analyzed using unpaired Student’s t-test. Data of H are analyzed using one-way ANOVA.

**Supplemental figure 5.**
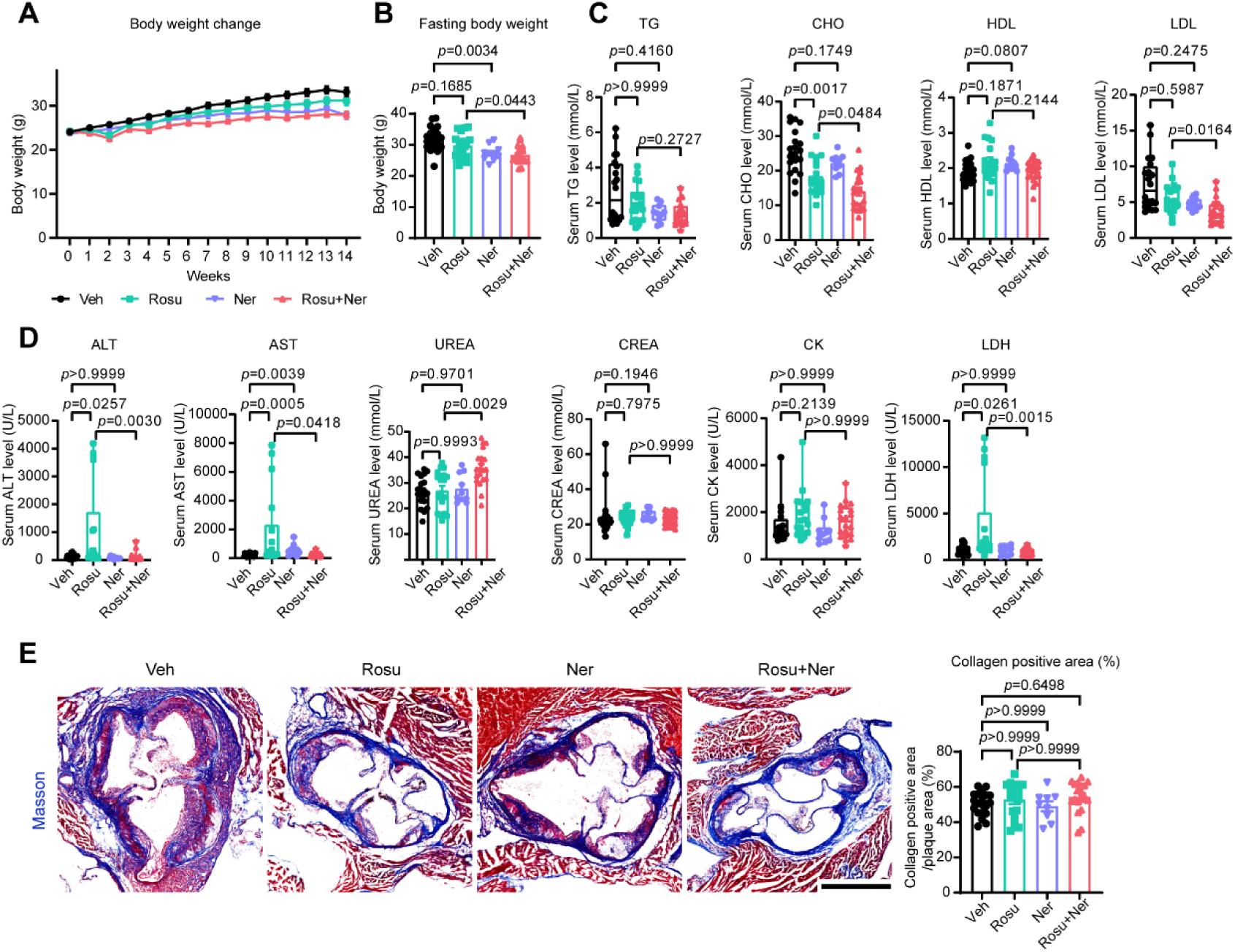
The effect of combination of Rosuvastatin and Neratinib on body weight, lipid profile, biochemistry parameters and lesional collagen content. Eight-week-old male *Ldlr*^−/−^ mice were randomly divided into four groups and orally administered 5 mg/kg Rosuvastatin, 20 mg/kg Neratinib or a combination of Rosuvastatin and Neratinib; n=10 Neratinib group, n=18 for Rosuvastatin group and n=20 for vehicle and combination group. (A) Body weight change of *Ldlr*^−/−^ mice. (B) Fasting body weight of each group. The mice were weighed after an overnight fasting for 12 h. (C and D) Serum lipid profile and biochemistry parameters of *Ldlr*^−/−^ mice. (E) Masson staining of aortic sinus cryosections showing the collagen content. Scale bar: 625 μm, n=9 for Neratinib group, n=17 for Rosuvastatin, n=20 for vehicle and n=19 for combination group. Abbreviations: ALT: alanine transaminase, AST: aspartate transaminase, CHO: total cholesterol, CK: creatine kinase, CREA: creatine, HDL: high-density lipoprotein, LDH: lactate dehydrogenase, LDL: low-density lipoprotein, Ner: Neratinib, TG: total triglyceride, Rosu: Rosuvastatin, Veh: vehicle. The data are presented as the means ± SEMs. Data of B and E were analysis using one-way ANOVA test. Data of TG and LDL in C, ALT, AST, CREA, CK, and LDH in D were analyzed using Kruskal-Wallis test. Data of CHO and HDL in C, UREA in D were analyzed using Brown-Forsythe and Welch ANOVA test.

## Supplemental tables

Supplemental table 1-9 are available in online data supplement as individual excel spreadsheet.

## References and Notes

1 Libby P. The changing landscape of atherosclerosis. Nature 2021; 592: 524–33.10.1038/s41586-021-03392-8.

2 Libby P. Inflammation and the pathogenesis of atherosclerosis. Vasc Pharmacol 2024; 154. 10.1016/j.vph.2023.107255.

3 Ridker PM, Everett BM, Thuren T, MacFadyen JG, Chang WH, Ballantyne C, et al. Antiinflammatory Therapy with Canakinumab for Atherosclerotic Disease. New England Journal of Medicine 2017; 377: 1119–31. 10.1056/NEJMoa1707914.

4 Kelly P, Lemmens R, Weimar C, Walsh C, Purroy F, Barber M, et al. Long-term colchicine for the prevention of vascular recurrent events in non-cardioembolic stroke (CONVINCE) : a randomised controlledtrial. Lancet 2024; 404: 125–33. 10.1016/S0140-6736(24)00968-1.

5 Ridker PM, MacFadyen JG, Everett BM, Libby P, Thuren T, Glynn RJ, et al. Relationship of C-reactive protein reduction to cardiovascular event reduction following treatment with canakinumab: a secondary analysis from the CANTOS randomised controlled trial. Lancet 2018; 391: 319–28. 10.1016/S0140-6736(17)32814-3.

6 Ridker PM, Devalaraja M, Baeres FMM, Engelmann MDM, Hovingh GK, Ivkovic M, et al. IL-6 inhibition with ziltivekimab in patients at high atherosclerotic risk (RESCUE): a double-blind, randomised, placebo-controlled, phase 2 trial. Lancet 2021; 397: 2060–9. 10.1016/S0140-6736(21)00520-1.

7 Pushpakom S, Iorio F, Eyers PA, Escott KJ, Hopper S, Wells A, et al. Drug repurposing: progress, challenges and recommendations. Nat Rev Drug Discov 2019; 18: 41–58. 10.1038/nrd.2018.168.

8 Verma S, Mazer CD, Connelly KA. Inflammation and cholesterol at the crossroads of vascular risk. Cell Metab 2023; 35: 1095–8. 10.1016/j.cmet.2023.06.011.

9 Subramanian A, Narayan R, Corsello SM, Peck DD, Natoli TE, Lu XD, et al. A Next Generation Connectivity Map: L1000 Platform and the First 1,000,000 Profiles. Cell 2017; 171: 1437-+. 10.1016/j.cell.2017.10.049.

10 Lamb J, Crawford ED, Peck D, Modell JW, Blat IC, Wrobel MJ, et al. The connectivity map: Using gene-expression signatures to connect small molecules, genes, and disease. Science 2006; 313: 1929–35. 10.1126/science.1132939.

11 LiverTox: Clinical and Research Information on Drug-Induced Liver Injury [Internet]. (Bethesda (MD): National Institute of Diabetes and Digestive and Kidney Diseases, 2012).

12 Stark AK, Davenport ECM, Patton DT, Scudamore CL, Vanhaesebroeck B, Veldhoen M, et al. Loss of Phosphatidylinositol 3-Kinase Activity in Regulatory T Cells Leads to Neuronal Inflammation. J Immunol 2020; 205: 78–89. 10.4049/jimmunol.2000043.

13 Pouwer MG, Pieterman EJ, Verschuren L, Caspers MPM, Kluft C, Garcia RA, et al. The BCR-ABL1 Inhibitors Imatinib and Ponatinib Decrease Plasma Cholesterol and Atherosclerosis, and Nilotinib and Ponatinib Activate Coagulation in a Translational Mouse Model. Frontiers in Cardiovascular Medicine 2018; 5. 10.3389/fcvm.2018.00055.

14 Lin ZM, Lin XC, Lai Y, Han CC, Fan XR, Tang J, et al. Ponatinib modulates the metabolic profile of obese mice by inhibiting adipose tissue macrophage inflammation. Front Pharmacol 2022; 13. 10.3389/fphar.2022.1040999.

15 Dutta B, Park JE, Kumar S, Hao PL, Gallart-Palau X, Serra A, et al. Monocyte adhesion to atherosclerotic matrix proteins is enhanced by Asn-Gly-Arg deamidation. Scientific Reports 2017; 7. 10.1038/s41598-017-06202-2.

16 Filik Y, Bauer K, Hadzijusufovic E, Haider P, Greiner G, Witzeneder N, et al. PI3-kinase inhibition as a strategy to suppress the leukemic stem cell niche in Ph plus chronic myeloid leukemia. Am J Cancer Res 2021; 11: 6042-+.

17 Paez-Mayorga J, Chen AL, Kotla S, Tao YT, Abe RJ, He ED, et al. Ponatinib Activates an Inflammatory Response in Endothelial Cells via ERK5 SUMOylation. Frontiers in Cardiovascular Medicine 2018; 5. 10.3389/fcvm.2018.00125.

18 Madonna R, Barachini S, Moscato S, Ippolito C, Mattii L, Lenzi C, et al. Sodium-glucose cotransporter type 2 inhibitors prevent ponatinib-induced endothelial senescence and disfunction: A potential rescue strategy. Vasc Pharmacol 2022; 142. 10.1016/j.vph.2021.106949.

19 Liu SYM, Tu HY, Wei XW, Yan HH, Dong XR, Cui JW, et al. First-line pyrotinib in advanced HER2-mutant non-small-cell lung cancer: a patient-centric phase 2 trial. Nat Med 2023; 29: 2079-+. 10.1038/s41591-023-02461-x.

20 Ma F, Yan M, Li W, Ouyang QC, Tong ZS, Teng YE, et al. Pyrotinib versus placebo in combination with trastuzumab and docetaxel as first line treatment in patients with HER2 positive metastatic breast cancer (PHILA): randomised, double blind, multicentre, phase 3 trial. Bmj-Brit Med J 2023; 383. 10.1136/bmj-2023-076065.

21 Saura C, Oliveira M, Feng YH, Dai MS, Chen SW, Hurvitz SA, et al. Neratinib Plus Capecitabine Versus Lapatinib Plus Capecitabine in HER2-Positive Metastatic Breast Cancer Previously Treated With ≥ 2 HER2-Directed Regimens: Phase III NALA Trial. J Clin Oncol 2020; 38: 3138-+. 10.1200/Jco.20.00147.

22 Bundred N, Porta N, Brunt AM, Cramer A, Hanby A, Shaaban AM, et al. Combined Perioperative Lapatinib and Trastuzumab in Early HER2-Positive Breast Cancer Identifies Early Responders: Randomized UK EPHOS-B Trial Long-Term Results. Clin Cancer Res 2022; 28: 1323–34. 10.1158/1078-0432.Ccr-21-3177.

23 Ichijo H, Nishida E, Irie K, tenDijke P, Saitoh M, Moriguchi T, et al. Induction of apoptosis by ASK1, a mammalian MAPKKK that activates SAPK/JNK and p38 signaling pathways. Science 1997; 275: 90–4. 10.1126/science.275.5296.90.

24 Lamb J. Innovation - The Connectivity Map: a new tool for biomedical research. Nat Rev Cancer 2007; 7: 54–60. 10.1038/nrc2044.

25 Deeks ED. Neratinib: First Global Approval. Drugs 2017; 77: 1695–704. 10.1007/s40265-017-0811-4.

26 Jhaveri K, Eli LD, Wildiers H, Hurvitz SA, Guerrero-Zotano A, Unni N, et al. Neratinib plus fulvestrant plus trastuzumab for HR-positive, HER2-negative, -mutant metastatic breast cancer: outcomes and biomarker analysis from the SUMMIT trial. Ann Oncol 2023; 34: 885–98. 10.1016/j.annonc.2023.08.003.

27 Barcenas CH, Hurvitz SA, Di Palma JA, Bose R, Chien AJ, Iannotti N, et al. Improved tolerability of neratinib in patients with HER2-positive early-stage breast cancer: the CONTROL trial. Ann Oncol 2020; 31: 1223–30. 10.1016/j.annonc.2020.05.012.

28 Dent P, Booth L, Roberts JL, Liu JC, Poklepovic A, Lalani AS, et al. Neratinib inhibits Hippo/YAP signaling, reduces mutant K-RAS expression, and kills pancreatic and blood cancer cells. Oncogene 2019; 38: 5890–904. 10.1038/s41388-019-0849-8.

29 Kheraldine H, Hassan AF, Alhussain H, Al-Thawadi H, Vranic S, Al Moustafa AE. Effects of neratinib on angiogenesis and the early stage of the embryo using chicken embryo as a model. Biomol Biomed 2024; 24: 575–81. 10.17305/bb.2023.9869.

30 Park YJ, An HT, Park JS, Park O, Duh AJ, Kim K, et al. Tyrosine kinase inhibitor neratinib attenuates liver fibrosis by targeting activated hepatic stellate cells. Scientific Reports 2020; 10. 10.1038/s41598-020-71688-2.

31 van Gerven J, Bonelli M. Commentary on the EMA Guideline on strategies to identify and mitigate risks for first-in-human and early clinical trials with investigational medicinal products. Brit J Clin Pharmaco 2018; 84: 1401–9. 10.1111/bcp.13550.

32 Tao G, Dagher F, Ghose R. Neratinib causes non-recoverable gut injury and reduces intestinal cytochrome P450 3A enzyme in mice. Toxicol Res-Uk 2022; 11: 184–94. 10.1093/toxres/tfab111.

33 Patel P, Naik MU, Naik U. Apoptosis Signal-Regulating Kinase (ASK1) Regulates Thrombosis in Part By Regulating cPLA phosphorylation-Dependent TxA Generation. Blood 2016; 128. 10.1182/blood.V128.22.3719.3719.

34 Rastogi S, Rizwani W, Joshi B, Kunigal S, Chellappan SP. TNF-α response of vascular endothelial and vascular smooth muscle cells involve differential utilization of ASK1 kinase and p73. Cell Death Differ 2012; 19: 274–83. 10.1038/cdd.2011.93.

35 Luyendyk JP, Piper JD, Tencati M, Reddy KV, Holscher T, Zhang R, et al. A novel class of antioxidants inhibit LPS induction of tissue factor by selective inhibition of the activation of ASK1 and MAP kinases. Arterioscl Throm Vas 2007; 27: 1857–63. 10.1161/Atvbaha.107.143552.

36 Yamawaki H, Pan S, Lee RT, Berk BC. Fluid shear stress inhibits vascular inflammation by decreasing thioredoxin-interacting protein in endothelial cells. J Clin Invest 2005; 115: 733–8. 10.1172/Jci200523001.

37 Ricci R, Sumara G, Sumara I, Rozenberg I, Kurrer M, Akhmedov A, et al. Requirement of JNK2 for scavenger receptor A-mediated foam cell formation in atherogenesis. Science 2004; 306: 1558–61. 10.1126/science.1101909.

38 Yamada S, Ding Y, Tanimoto A, Wang KY, Guo X, Li Z, et al. Apoptosis Signal-Regulating Kinase 1 Deficiency Accelerates Hyperlipidemia-Induced Atheromatous Plaques via Suppression of Macrophage Apoptosis. Arterioscl Throm Vas 2011; 31: 1555–U172. 10.1161/Atvbaha.111.227140.

39 Liu QL, Pan JL, Bao LR, Xu CX, Qi Y, Jiang B, et al. Major Vault Protein Prevents Atherosclerotic Plaque Destabilization by Suppressing Macrophage ASK1-JNK Signaling. Arterioscl Throm Vas 2022; 42: 580–96. 10.1161/Atvbaha.121.316662.

40 Al Ghouleh I, Frazziano G, Rodriguez AI, Csányi G, Maniar S, St Croix CM, et al. Aquaporin 1, Nox1, and Ask1 mediate oxidant-induced smooth muscle cell hypertrophy. Cardiovasc Res 2013; 97: 134–42. 10.1093/cvr/cvs295.

41 Tasaki T, Yamada S, Guo X, Tanimoto A, Wang KY, Nabeshima A, et al. Apoptosis Signal-Regulating Kinase 1 Deficiency Attenuates Vascular Injury-Induced Neointimal Hyperplasia by Suppressing Apoptosis in Smooth Muscle Cells. Am J Pathol 2013; 182: 597–609. 10.1016/j.ajpath.2012.10.008.

42 Izumi Y, Kim S, Yoshiyama M, Izumiya Y, Yoshida K, Matsuzawa A, et al. Activation of apoptosis signal-regulating kinase 1 in injured artery and its critical role in neointimal hyperplasia. Circulation 2003; 108: 2812–8. 10.1161/01.Cir.0000096486.01652.Fc.

43 Hang LW, Peng Y, Xiang R, Li XD, Li ZL. Ox-LDL Causes Endothelial Cell Injury Through ASK1/NLRP3-Mediated Inflammasome Activation via Endoplasmic Reticulum Stress. Drug Des Dev Ther 2020; 14: 731–43. 10.2147/Dddt.S231916.

44 Zhang JL, Du BB, Zhang DH, Li H, Kong LY, Fan GJ, et al. OTUB1 alleviates NASH through inhibition of the TRAF6-ASK1 signaling pathways. Hepatology 2022; 75: 1218–34. 10.1002/hep.32179.

45 Feng H, Cao JL, Zhang GY, Wang YG. Kaempferol Attenuates Cardiac Hypertrophy via Regulation of ASK1/MAPK Signaling Pathway and Oxidative Stress. Planta Med 2017; 83: 837–45. 10.1055/s-0043-103415.

46 Ho KL, Ramin C, Shing JZ, Taparra K, Vo JB. Effect of county-level income and rurality on cardiovascular disease mortality among Asian and Pacific Islander breast cancer survivors in the US, 2000-2018. Cancer Res 2023; 84. 10.1158/1538-7445.Am2024-4828.

47 Pudil R, Danzig V, Vesely J, Málek F, Táborsky M, Elbl L, et al. 2022 ESC Guidelines on cardio-oncology developed in collaboration with the European Hematology Association (EHA), the European Society for Therapeutic Radiology and Oncology (ESTRO) and the International Cardio-Oncology Society (IC-OS). Cor Vasa 2023; 65: 350–434. 10.33678/cor.2023.032.

48 Lenihan D, Suter T, Brammer M, Neate C, Ross G, Baselga J. Pooled analysis of cardiac safety in patients with cancer treated with pertuzumab (vol 23, pg 791, 2012). Ann Oncol 2019; 30: 1021-. 10.1093/annonc/mdy533.

49 von Minckwitz G, Procter M, de Azambuja E, Zardavas D, Benyunes M, Viale G, et al. Adjuvant Pertuzumab and Trastuzumab in Early HER2-Positive Breast Cancer. New England Journal of Medicine 2017; 377: 122–31. 10.1056/NEJMoa1703643.

50 Cameron D, Brown J, Dent R, Jackisch C, Mackey J, Pivot X, et al. Adjuvant bevacizumab-containing therapy in triple-negative breast cancer (BEATRICE): primary results of a randomised, phase 3 trial. Lancet Oncol 2013; 14: 933–42. 10.1016/S1470-2045(13)70335-8.

## Supplemental references

51 Xu MY, Xu JJ, Kang LJ, Liu ZH, Su MM, Zhao WQ, et al. Urolithin A promotes atherosclerotic plaque stability by limiting inflammation and hypercholesteremia in Apolipoprotein E-deficient mice. Acta Pharmacol Sin 2024. 10.1038/s41401-024-01317-5.

52 Subramanian A, Narayan R, Corsello SM, Peck DD, Natoli TE, Lu XD, et al. A Next Generation Connectivity Map: L1000 Platform and the First 1,000,000 Profiles. Cell 2017; 171: 1437-+. 10.1016/j.cell.2017.10.049.

53 Andueza A, Kumar S, Kim J, Kang DW, Mumme HL, Perez JI, et al. Endothelial Reprogramming by Disturbed Flow Revealed by Single-Cell RNA and Chromatin Accessibility Study. Cell Rep 2020; 33. 10.1016/j.celrep.2020.108491.

54 Alsaigh T, Evans D, Frankel D, Torkamani A. Decoding the transcriptome of calcified atherosclerotic plaque at single-cell resolution. Commun Biol 2022; 5. 10.1038/s42003-022-04056-7.

55 Wang ZH, Wang MM, Zhang MX, Xu KK, Zhang XS, Xie Y, et al. High-affinity SOAT1 ligands remodeled cholesterol metabolism program to inhibit tumor growth. Bmc Med 2022; 20. 10.1186/s12916-022-02436-8.

56 Wang ZH, Wu WB, Guan XC, Guo S, Li CW, Niu RX, et al. 20(S)-Protopanaxatriol promotes the binding of P53 and DNA to regulate the antitumor network multiomic analysis. Acta Pharmaceutica Sinica B 2020; 10: 1020–35. 10.1016/j.apsb.2020.01.017.

57 Molina DM, Jafari R, Ignatushchenko M, Seki T, Larsson EA, Dan C, et al. Monitoring Drug Target Engagement in Cells and Tissues Using the Cellular Thermal Shift Assay. Science 2013; 341: 84–7. 10.1126/science.1233606.

58 Dal Piaz F, Saltos MBV, Franceschelli S, Forte G, Marzocco S, Tuccinard T, et al. Drug Affinity Responsive Target Stability (DARTS) Identifies Laurifolioside as a New Clathrin Heavy Chain Modulator. J Nat Prod 2016; 79: 2681–92. 10.1021/acs.jnatprod.6b00627.

